# Spatial cumulant models enable spatially informed treatment strategies and analysis of local interactions in cancer systems

**DOI:** 10.1101/2022.05.07.491050

**Authors:** Sara Hamis, Panu Somervuo, J. Arvid Ågren, Dagim Shiferaw Tadele, Juha Kesseli, Jacob G. Scott, Matti Nykter, Philip Gerlee, Dmitri Finkelshtein, Otso Ovaskainen

## Abstract

Theoretical and applied cancer studies that use individual-based models (IBMs) have been limited by the lack of a mathematical formulation that enables rigorous analysis of these models. However, spatial cumulant models (SCMs), which have arisen from theoretical ecology, describe population dynamics generated by a specific family of IBMs, namely spatio-temporal point processes (STPPs). SCMs are spatially resolved population models formulated by a system of differential equations that approximate the dynamics of two STPP-generated summary statistics: first-order spatial cumulants (densities), and second-order spatial cumulants (spatial covariances).

We exemplify how SCMs can be used in mathematical oncology by modelling theoretical cancer cell populations comprising interacting growth factor-producing and non-producing cells. To formulate model equations, we use computational tools that enable the generation of STPPs, SCMs and mean-field population models (MFPMs) from user-defined model descriptions (Cornell et al., 2019). To calculate and compare STPP, SCM and MFPM-generated summary statistics, we develop an application-agnostic computational pipeline. Our results demonstrate that SCMs can capture STPP-generated population density dynamics, even when MFPMs fail to do so. From both MFPM and SCM equations, we derive treatment-induced death rates required to achieve non-growing cell populations. When testing these treatment strategies in STPP-generated cell populations, our results demonstrate that SCM-informed strategies outperform MFPM-informed strategies in terms of inhibiting population growths. We thus demonstrate that SCMs provide a new framework in which to study cell-cell interactions, and can be used to describe and perturb STPP-generated cell population dynamics. We, therefore, argue that SCMs can be used to increase IBMs’ applicability in cancer research.

**Statements and Declarations:** The authors have no competing interests to declare that are relevant to the content of this article.

## 1 Introduction

All biological systems are composed of interacting parts. In cancer systems, cells can interact with each other by, for example, exchanging signalling molecules and competing for resources such as space and nutrients. Such cell-cell interactions have been identified as factors that drive eco-evolutionary dynamics of cancer cell populations (Merlo et al., 2006; Reynolds et al., 2020). Consequently, these interactions have been proposed as cancer treatment targets, where the general premise is that treatments can perturb cellcell interactions and, by extension, disease trajectories. Such perturbations can lead to reduced tumour growths and disease burdens (Dominiak et al., 2020; Brücher and Jamall, 2014; West and Newton, 2019). Cell-cell interactions, and the impact that they have on cancer system dynamics, have been mathematically studied by researchers using a variety of mathematical models including evolutionary game theory (EGT) models (Kaznatcheev et al., 2019; Archetti et al., 2015; Farrokhian et al., 2022), other ordinary differential equation (ODE) models (Hillen et al., 2013; Poleszczuk and Enderling, 2018; Jenner et al., 2018), partial differential equation (PDE) models (Haridas et al., 2017; Lorenzi et al., 2017; Hinow et al., 2009), and individual-based models (IMBs) (Stichel et al., 2017; Campenni et al., 2020; Hamis et al., 2021).

In IBMs, cells can be modelled as individuals that evolve on a spatial domain and partake in interactions with other cells in their neighbourhood (Metzcar et al., 2019; Chamseddine and Rejniak, 2020). IBMs are naturally able to incorporate local cell-cell interactions, spatial population structures and stochastic variations between cells and biological events. However, theoretical and experimental studies that use IBMs have been limited by the lack of a mathematical formulation that enables rigorous analyses of these models. On the other hand, in mean-field population models (MFPMs), such as ODE and EGT models, the density dynamics of cancer cell subpopulations can be described by a set of mathematically tractable ODEs after imposing a set of simplifying modelling assumptions. In such models, cells are not considered as individuals but, instead, they are collectively viewed as part of a population or subpopulation. With their analytical and mathematically tractable formulation, MFPMs have been used to quantify cell-cell interactions in *in vitro* experiments (Kaznatcheev et al., 2019), and are currently being used to inform personalised treatment strategies in a clinical prostate cancer trial (Zhang et al., 2017) (see also matters arising in response to the reporting of this study (Mistry, 2021)). However, MFPMs notably assume that the modelled system is “well-mixed” so that each cell interacts with all other cells in the system with equal probability. Therefore, MFPMs can not faithfully capture the dynamics of spatially structured systems with localised cell-cell interactions, such as solid tumours (Waclaw et al., 2015). In fact, previous mathematical modelling work has shown that imposing spatial constraints on interactions between individuals, via IBMs, often results in system dynamics that vastly contradict that simulated by MFPMs (Durrett and Levin, 1994).

Due to the pre-clinical and clinical applications of mathematical oncology models (Rockne et al., 2019), there exists a need to formulate models that capture localised cell-cell interactions, maintain cell discreteness, and are both spatio-temporally resolved and mathematically tractable. In this article, we describe an approach to achieve this by using spatial cumulant models (SCMs) which, to our knowledge, have not previously been applied to study cancer cell systems. SCMs are spatially resolved population models that are translated from a specific family of IBMs, namely spatio-temporal point processes (STPPs). Following a mathematical manipulation that involves a perturbation expansion around mean-field equations in the limit of long-ranged interactions (as explained in Section 2), SCMs approximate two STPP-generated summary statistics: first-order spatial cumulants (densities) and second-order spatial cumulants (spatial covariances). If we let 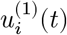 denote the density of subpopulation *i*, and 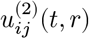 denote the spatial covariance of individuals (here cells) from subpopulations *i* and *j*, then SCMs dictate that

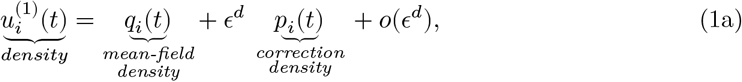

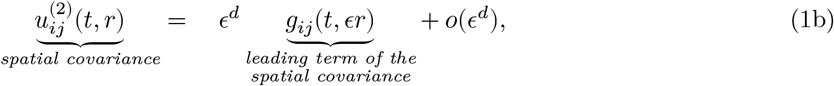

where *t* is the independent time variable and *r* denotes pair-wise distances between individuals. In Eq.1a, *q*_*i*_(*t*) is the density as computed by MFPMs and *p*_*i*_(*t*) is a correction term that depends on (1) the MFPM densities *q*_*i*_(*t*), (2) the leading spatial covariance terms *g*_*ij*_(*t, ϵr*), and (3) all corrections *p*_*i*_(*t*), where *i, j* = 1, .., *S*, and *S* denotes the number of subpopulations in the modelled system. Thus the correction terms account for the spatial structure of the population via their dependence on *g*_*ij*_(*t, ϵr*). The modelled individuals inhabit a *d*-dimensional spatial domain, and the scaling factor *ϵ* = 1/ℓ, ℓ > 0, results from the perturbation expansion, in which ℓ is an interaction length scale. Thus, SCM densities reduce to MFPM densities when ℓ → ∞. The terms *o*(*ϵ*^*d*^) can be omitted in practical applications as they go to zero faster than does *ϵ*^*d*^.

Since Eqs.1a,b are formulated in the limit of long-ranged interactions it follows that, in the context of cancer research, SCMs are appropriate to use when studying cell-cell interactions that extend beyond only immediately neighbouring cells. Such interactions include communication via diffusible substrates and resource competition. Formulating the right-hand sides of Eqs.1a,b for a specific biological problem can be a mathematically cumbersome task. However, Cornell *et al*. (2019) developed a mathematical framework and a computational software that enables the generation of rate equations for the quantities *q*_*i*_*(t), p*_*i*_*(t)* and *g*_*ij*_*(t, ϵr)* for a large family of biological processes. More specifically, the software enables the generation of (1) STPPs and (2) MFPM and SCM rate equations for any user-defined biological system that can be described by one or more reactant-catalyst-product (RCP) processes. An RCP-processes is formulated by a set of reactants (points that are consumed in the process), catalysts (points that catalyse the process without being consumed in it), and products (points that are produced in the process). In this study, these points represent the center of mass of cells. Cornell et al.’s framework thus unifies a class of IBMs, *i*.*e*., STPPs, with a class of analytical and spatially structured population models, *i*.*e*., SCMs. Accordingly, we refer to their software as the `Unified Framework (UF) Software` throughout this article.

SCMs were first presented as a rigorous mathematical framework by Ovaskainen *et al*. (2014). The models build upon earlier studies of spatial moment equations in theoretical ecology (Ovaskainen and Cornell, 2006; Bolker and Pacala, 1997; Law et al., 2003) and probability theory (Finkelshtein et al., 2009; Kondratiev et al., 2008), and on probability theorists’ work on STPPs, also known as “Markov evolutions in the space of locally finite configurations”, (Kondratiev and Skorokhod, 2011; Finkelshtein et al., 2011). Spatial cumulants are closely related to spatial moments (Section 2.4), and we remark that spatial moment techniques have been used to derive analytical, deterministic approximations for cell population dynamics generated by different types of IBMs. These include on-lattice models in which cells are restricted to inhabit pre-defined lattice points, that can be occupied by one or zero cells (Baker and Simpson, 2010; Markham et al., 2015) and STPPs (Binny et al., 2016; Surendran et al., 2018). On-lattice IBMs explicitly account for volume exclusion, as opposed to STPPs which do not. STPPs instead generate points that can represent the center of mass of deformable cells that can move in continuous space and interact with their environment via probabilistic kernels. This brief discussion exemplifies that all IBMs are associated with different caveats and benefits. Therefore, a bio-mathematical modeller’s choice of IBM and, by extension, options for describing IBM dynamics analytically, should be considered on an application-specific basis. The aforementioned cell population studies that used spatial moment techniques all challenged the view that there exists a dichotomy between individual-based and analytical models. We conjecture that one of the reasons that such techniques are not more widely used in cell biology research is that the relevant model equations have only been derived for specific cell processes, as opposed to general processes. Accordingly, a modeller wanting to use spatial moment techniques for describing the dynamics of cell populations, in which specific cell actions and interactions occur, might face a mathematically technical and time-consuming task of deriving spatial moment (or cumulant) equations with appropriate closure and approximation methods. The `UF-Software` tackles this issue of lack-of-generality by allowing a modeller to formulate spatial cumulant equations for any RCP-process. Moreover, the pipeline codes developed in this study enable a modeller to easily implement and numerically solve SCM equations. These pipeline codes also enable straightforward comparisons between STPP, SCM and MFPM-generated population dynamics.

The remainder of this article is structured as follows: In Section 2, we provide an introductory description of the mathematical theory that underlies SCMs and the `UF-Software`. In Section 3.1, we propose a practical step-by-step pipeline for using the `UF-Software` to implement STPPs, MFMPs and SCMs. Thereafter, as proof-of-concept, we exemplify how SCMs and the `UF-Software` can be used in mathematical cancer research by modelling dynamic, spatially structured cancer cell populations in which cells interact via diffusible substrates and are subjected to drugs (Sections 3.2-4.2). We demonstrate that SCMs can be used to both describe and perturb density dynamics of simulated cell populations generated by STPPs.

## 2 Mathematical theory of spatial cumulant models

Sections 2.1-2.10 are intended to comprise an introductory and conceptually self-contained background to SCMs. For more rigorous details of this mathematical theory, we refer readers to previous works (Ovaskainen and Cornell, 2006; Ovaskainen et al., 2014; Cornell et al., 2019). For ease of presentation we, throughout this article, consider spatial domains that are two-dimensional and have periodic boundary conditions. Moreover, we consider systems that are translationally invariant, meaning that pair-wise distances between cells influence system dynamics, whereas absolute positions of cells do not. This is equivalent to modelling a space-homogeneous system that lacks background variations, such as a typical *in vitro* system. Note, however, that the `UF-Software` can be modified to handle one, two, and threedimensional spatial domains, as well as space-heterogeneous systems (Cornell et al., 2019).

### 2.1 Spatio-temporal point patterns and spatio-temporal point processes

Consider a cell population that evolves in time and space. A snapshot of the population can be represented by a point pattern in which the center of mass of the *n*th cell is represented by a point *x*_*n*_ ∈ *D*, where *D* ⊂ ℝ^*d*^ denotes the spatial domain that the population inhabits. In this article, we set *D* to be a plane spanning two spatial dimensions, so that *d* = 2 and 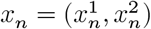. A point pattern comprising *N*_t_ points at time *t* can thus be described by a set of points *η*_*t*_ in continuous space such that

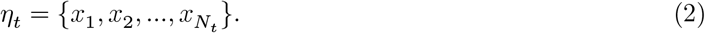

In order to categorise the individuals in the population, each point in *η*_*t*_ can further be coupled with a descriptive mark. Here, we let the mark *m*_*n*_ denote the subpopulation to which the individual in position *x*_*n*_ belongs so that *m*_*n*_ can take the value 1, 2, …, *S*, where *S* denotes the number of subpopulations in the modelled system. When marks are included, a point pattern describing a population snapshot can thus be described by a set of marked points

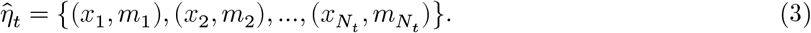

Two summary statistics that can be used to describe point patterns are densities and spatial covariances. Instructions on how to calculate these quantities from static point patterns are provided in Section 2.9. A point pattern that evolves in time and space is called a spatio-temporal point pattern, and an STPP is a mathematical model that can generate spatio-temporal point patterns in continuous time and continuous space. In this article, we will denote spatio-temporal point patterns by *η*, whether they are marked or not. Intuitively, an STPP can be thought of as a collection of stochastic processes that makes points appear or disappear from *η*. In the context of cell biology, such processes can, for example, describe cell division, cell movement, and cell death.

### 2.2 A verbal description of the relationship between spatio-temporal point processes and spatial cumulant models

The STPP rules that govern the evolution of spatio-temporal point patterns are of a stochastic nature. Therefore, a general STPP can generate an ensemble of different realisations, *i*.*e*., spatio-temporal point patterns, from a given initial cell configuration. It is, in general, impossible to predict how one specific realisation will evolve in time and space. However, if we generate multiple STPP realisations, then SCMs can be used to deterministically approximate how densities and spatial covariances are expected to evolve on average for the realisations.

### 2.3 Reactant-catalyst-product processes and interaction kernels

The `UF-Software` can be used to generate STPPs, MFPMs and SCMs for any biological system that can be described by one or more of RCP-processes. One-point RCP-processes include density-independent cell death, as the death of a cell in location *x*_*V*_ with mark *m* can be described by a single reactant, *i*.*e*., point, (*x*_*V*_, *m*). Such one-point RCP-processes are associated with a scalar model parameter that describes the rate (probability per time unit) at which the process occurs. Multiple-point RCP-processes include density-independent cell division and cell movement. More precisely, density-independent cell division can be described as an RCP-process in which a cell of mark *m* in location *x*_*V*_ produces a daughter cell with the same mark in location *x*_*w*_, so that point (*x*_*V*_,*m*) is a catalyst and point (*x*_*w*_,*m*) is a product. Similarly, if a cell in location *x*_*V*_ with mark *m* moves to location *x*_*u*_, then point (*x*_*V*_,*m*) is a reactant and point (*x*_*u*_,*m*) is a product. The rate at which such multiple-point RCP-processes occur are described by interaction kernels that depend on pair-wise distances between points.

The `UF-Software` includes two pre-defined, radially symmetric, interaction kernels, being the Gaussian kernel *a*_G_ and the Top-Hat kernel *a*_TH_ such that, on a two-dimensional spatial domain,

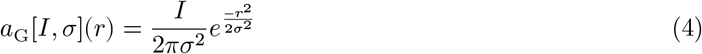

and

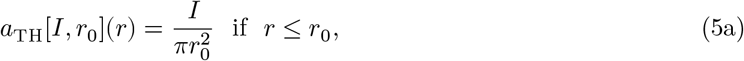

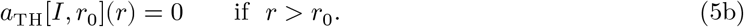

Here *I* denotes the total integral of the interaction kernel, *σ* is one kernel standard deviation, *r*_0_ is a kernel interaction radius, and *r* is the distance between two interacting points, *i*.*e*., reactants, catalysts or products. Note that the `UF-Software` can be modified to include any user-defined, non-negative and integrable interaction kernels. These kernels need not be maximised for zero cell-pair separations.

### 2.4 The relationship between spatial moments and spatial cumulants

The spatial cumulants are closely related to the spatial moments, which are also called correlation functions. The first spatial moment, *k*^(1)^(*x*), measures the expected density of individuals at location *x*. It follows that the expected number of individuals in an area Λ at a time point *t* can be calculated by

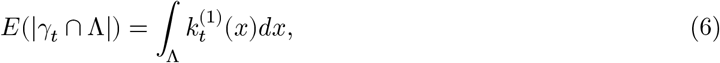

where *γ*_*t*_ is a point configuration. The second spatial moment, *k*^(2)^(*x*_1_, *x*_2_), can similarly be used to calculate the product of ‘the number of individuals in area Λ_1_’ and ‘the number of individuals in area Λ_2_’, such that

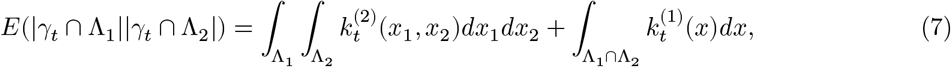

where the last term has been included in order to handle self-pairs (Ovaskainen et al., 2020). The first and second-order spatial cumulants are respectively denoted by 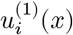 and 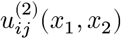, where

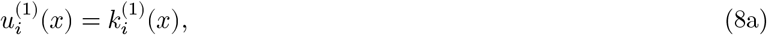

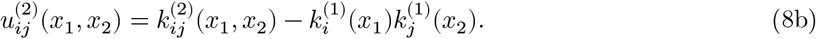

Here, the subscripts *i* and *j* have been included to denote different subpopulations in the modelled system, so that *i, j* = 1, .., *S*, where *S* is the number of modelled subpopulations.

From Eqs.8a,b we can see that the first and second-order spatial moments and cumulants carry the same information. To then understand why the spatial cumulants have been introduced, note first that spatial moments can be defined up to any order, and that dynamical systems of spatial moments generally are unclosed. For example, in systems in which pair-wise interactions occur, the dynamics of the *v*th-order spatial moment depends on the *v* + 1st-order spatial moment. Therefore, equations that describe spatial moment dynamics are generally impossible to solve without enforcing some sort of truncation or approximation closure scheme. The same closure problem arises for equations that describe the dynamics of spatial cumulants. However, in the limit of long-ranged interactions (Sections 2.6, 2.7), the spatial cumulant formulation enables the derivation of exact equations in closed form that describe first and second-order spatial cumulant dynamics of STPP-generated populations governed by RCP-processes (Section 2.8).

### 2.5 The relationship between Markov operators and operators for spatial cumulant rate equations

If we let *F* denote a real-valued function that describes some quantity of an STPP-generated realisation *γ*, and *µ* denotes a probability measure on Γ, then

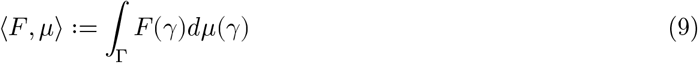

describes the value of *F* averaged over all realisations *γ* ∈ Γ. The rate of change of the pairing *⟨F, µ⟩* can be obtained by

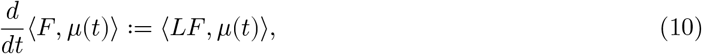

where *L* is Markov operator, that acts on *F* and describes a process that causes points to appear on, or disappear from, a set of points *η* (Eq.2). In order to describe a cell population that is governed by *B* biological processes (here, RCP-processes), the system’s total Markov operator can be written as a linear combination of Markov operators *L*_*b*_ such that

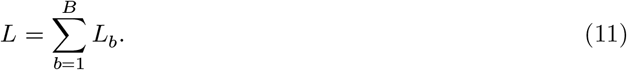

For any such Markov operator *L*, there exists a corresponding operator *L*^Δ^ that describes the time evolution of spatial moments of any order such that

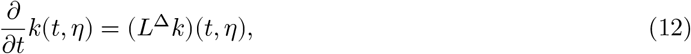

where *L*^Δ^ is a linear operator that depends on interaction kernels of the form *a*(*x*) (Section 2.3). Similarly, a Markov operator *L* be mapped to an operator *Q*^Δ^ that describes the time evolution of spatial cumulants of any order such that

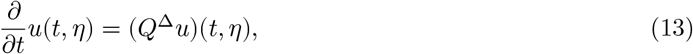

where *Q*^Δ^ is a sum of the linear term *L*^Δ^ and a non-linear term *M*^Δ^,

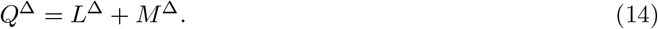

Deriving the operators *L*^Δ^ and *Q*^Δ^ from a Markov operator *L* can be a mathematically challenging and laborious task. However, for any system of RCP-processes, the `UF-Software` can handle the formulation of *L, L*^Δ^ and *Q*^Δ^ “under-the-hood”.

### 2.6 The perturbation expansion

As a first step towards circumventing the problem of moment closure in population-dynamics problems, Ovaskainen and Cornell (2006) initially proposed using a heuristic perturbation expansion in which interactions between points (individuals) were considered to be long-ranged, but not global. Note that Markov operators *L* (Eq.11) generally can be described using interaction kernels *a*(*x*) (Section 2.3). However, in the limit of long-ranged interactions, Ovaskainen *et al*. (2006) introduced scaled Markov operators *L*_*ϵ*_ that are described by scaled interaction kernels *a*_*ϵ*_*(x)*, such that

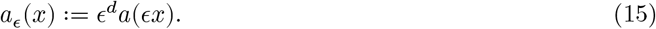

The integrals of *a(x)* and *a*_*ϵ*_*(x)* over ℝ^*d*^ have the same value, but when *ϵ → 0*, the scaled interaction kernels become increasingly long-ranged and flat so that each individual interacts with an increasing number of other individuals. In the perturbation expansion proposed by Ovaskainen and Cornell (2006), population densities *k*^*(1)*^*(t)* were expressed as

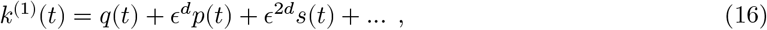

where *ϵ =* 1*/ℓ* and ℓ is a characteristic interaction length-scale. In Eq.16, the term *q*(*t*) denotes the density as calculated by MFPMs, and the higher-order terms *p(t), s(t)*, … are correction terms that account for the spatial structure of the modelled population. The perturbation expansion in Eq.16 has two notable practical advantages: (1) the calculated density *k*^(1)^*(t)* tends to the MFPM density when *ϵ →* 0, and (2) if we calculate *k*^(1)^*(t)* with correction terms up to the *n*th order, then we know beforehand that the error will behave as *o(ϵ*^*nd*^). The correction terms *p, s*, … in Eq.16 can in principle be derived up to arbitrarily high orders, but truncating the perturbation expansion after one correction terms yields

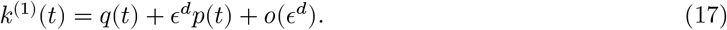

Ovaskainen and Cornell 2006 derived the expressions for *q(t)* and *p(t)* for a specific biological system and, based on their mathematical results, they argued that including only one correction term in the density calculation (Eq.17) is expected to suffice to capture STPP dynamics for many practical applications. Building on the heuristic work presented in 2006, Ovaskainen *et al*. later (2014) developed a rigorous mathematical framework for deriving spatial moment and cumulant rate equations from Markov operators, where the authors expected that spatial cumulants (of any orders) could be expressed as

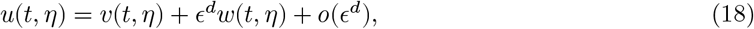

where *v*(*t, η*) is a mean-field term and *w*(*t, η*) is a first-order correction term. The authors showed that, for a specific model system (the spatial and stochastic logistic model) higher-order correction terms in Eq.18 can be omitted in the limit of long-ranged interactions. Making this result more generalisable, in 2019, Cornell *et al*. presented the `UF-Software` which enables the automatic generation of *v*(*t, η*) and *w*(*t, η*) for the first and second-order spatial cumulants in the limit of long-ranged interactions for any system that is formulated in terms of RCP-processes. Note that, in this study, only the number of points (or densities) and pair-wise distances between points in *η* are used in calculations, whereas other information regarding *η* is omitted. Therefore, dropping the explicit *η*-dependency in Eq.18, we may write the first and second-order spatial cumulants as *u*^(1)^(*t*) and *u*^(2)^(*t, r*) respectively (Eq.1a,b).

### 2.7 The model rescaling

Recall that the spatial moment dynamics generated by a Markov operator *L* can be described by an operator *L*^Δ^ (Eq.12). Such a Markov operator *L* generally depends on interaction kernels of the form *a*(*x*) (Section 2.3). If we, instead, consider a Markov operator *L*_*ϵ*_ that is described by the same (RCP) processes as *L*, but uses scaled interaction kernels of the form *a*_*ϵ*_(*x*) (Eq.15), we must introduce a new, scaled operator 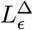 to describe the spatial moment dynamics generated by *L*. Thus, if we let 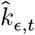 denote spatial moments that are generated by an STPP with scaled interaction kernels *a*_*ϵ*_(*x*), then

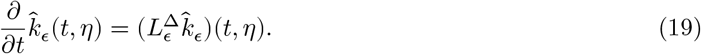

Note that Eq.19 is unclosed for general Markov operators *L*_*ϵ*_. In order to describe STPP-generated population dynamics by a system of closed equations, Ovaskainen *et al*. (2014) used a model-rescaling approach. To choose an appropriate rescaling, the authors remarked that the model should satisfy two important properties in order to be practical and relevant for biological applications: (1) when the interaction length *ℓ* goes to infinity, the model dynamics should tend to MFPM dynamics, as each individual would interact with an increasing number of other individuals. (2) The spatial clustering of the individuals should scale with *ℓ, i*.*e*., the model should scale with respect to space. The authors therefore introduced a scaling operator *S*_*c*_ that rescales a point configuration *η* such that

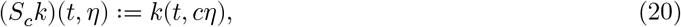

where

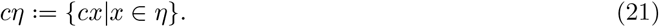

A renormalised version of the operator 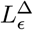 can then be defined as

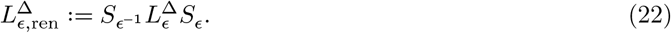

If we let *k*_*ϵ*_(*t, η*) denote the solution to the spatial moment equation that corresponds to this renormalised operator 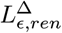, then

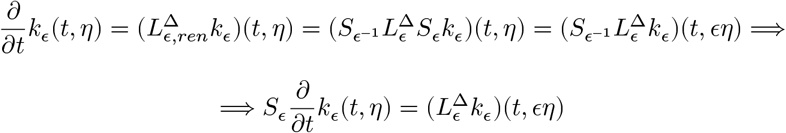

and thus,

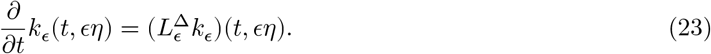

Note that the spatial moments are scaled in different ways in Eqs. 19 and 23. Together, these two equations enable the comparison of simulation-based results and analytical solutions. In other words, 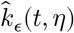 (which is a simulation-generated estimate of the spatial moment) and *k*_*ϵ*_(*t, ϵη*) (which is an analytical expression of the spatial moment) should agree. Therefore,

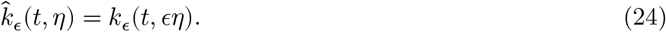

Analogously to Eq.23, rate equations for scaled spatial cumulants take the form

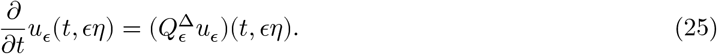

The assumed form of the spatial cumulants in the limit of long-ranged interactions (Eq.18) can be substituted into Eq.25. After such a substitution, the dynamical system is still unclosed so that the dynamics of the n-th order spatial cumulant *u*^(*n*)^(*t, ϵη*) depends on higher-order spatial cumulants. However, when solving for *v*(*t, η*) and *w*(*t, η*) (Eq.18), algebraic manipulation will show that *u*^(*n*)^(*t, ϵη*) = *o*(*ϵ*^*d*^) for *n* > 2, and thus can be omitted. This means that the model-rescaling approach (Section 2.7) together with the perturbation expansion (Section 2.6) enables the generation of closed, exact equations for the dynamics of the first and second-order spatial cumulants generated by STPPs.

The `UF-Software` can generate analytical expressions for *v*(*t, η*) and *w*(*t, η*) for the first and secondorder spatial cumulants from any model description formulated by one or more RCP-processes. Thus, when using the `UF-Software`, a user does not need to derive any operators, or perform any algebraic calculations to obtain mathematical expressions for *v*(*t, η*) and *w*(*t, η*).

### 2.8 Spatial cumulant model equations

The `UF-Software` formulates *v*(*t, η*) and *w*(*t, η*) in terms of rate equations for three auxiliary functions: *q*_*i*_(*t*), *p*_*i*_(*t*) and *g*_*ij*_(*t, ϵr*) which are used to formulate the first and second-order spatial cumulants. By comparing Eqs.1 and 18 we can see that, for the first-order spatial cumulant, *v*(*t, η*) = *q*_*i*_(*t*) and *w*(*t, η*) = *p*_*i*_(*t*). And, for the second-order spatial cumulant, *v*(*t, η*) = 0 and *w*(*t, η*) = *g*_*ij*_(*t, ϵr*). For all subpopulations *i, j* = 1, 2, .., *S*, the `UF-Software` generates the right-hand sides of the differential equations

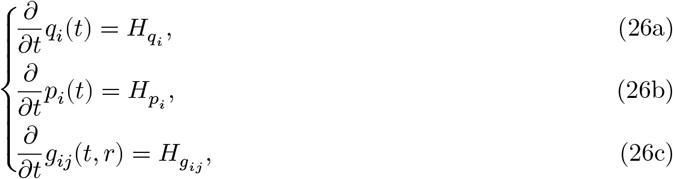

where 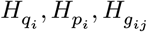 are exact expressions (Cornell et al., 2019). The expressions 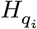 generally depend on all subpopulation densities *q*_1_(*t*), …, *q*_*S*_(*t*). Similarly, the expressions 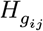 generally depend on all densities and leading spatial covariance terms *g*_*ij*_(*t, ϵr*). Lastly, the correction expressions 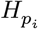 depend on all densities, leading spatial covariance terms and correction terms *p*_*i*_(*t*). generally

In order to calculate SCMs in practice, we first use the `UF-Software` to generate the the expressions for 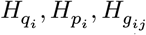. Thereafter, we use the pipeline codes developed for this study (Section 3.1) to numerically calculate *q*_*i*_(*t*), *g*_*ij*_(*t, ϵr*) and *p*_*i*_(*t*), in that order. By inserting these numerical results into Eqs.1a,b, densities and spatial covariances can be obtained. Note that, for many RCP-processes, the expressions for 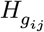 include convolution integrals, which are easier to work with in Fourier space than in real space. Therefore, `UF-Software`-generated expressions for 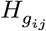 generally include Fourier transforms of interaction kernels and variables *g*_*ij*_(*t, ϵr*).

### 2.9 Calculating spatial moments and cumulants from point patterns

In the space-homogenous case, the static first-order spatial moment, 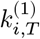, and cumulant, 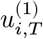, of subpopulation *i* at time *T* can be directly estimated from a point pattern as

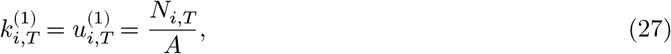

where *N*_*i,T*_ denotes the number of type *i* individuals on domain *D* with area *A*. Static second-order spatial moments at time *T* can be approximated from a point pattern by

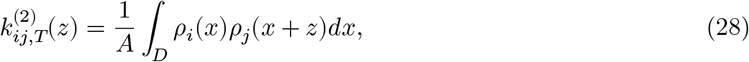

where *ρ*_*i*_ denotes the density of subpopulation *i*, and *z* is a spatial displacement so that *z* = (*z*^1^, *z*^2^) when *D* is two-dimensional (Dieckmann et al., 2000). The pair densities 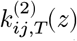 are also called auto-correlations (for *i* = *j*) and cross-correlations (for *i* ≠ *j*). When approximating_*ij*,_s_*T*_tatic second-order spatial moments for space-homogenous systems, the magnitude *r* = |*z*| of the displacement in Eq.28 is of importance, whereas the direction of *z* is not. In practical applications, we can thus construct a vector

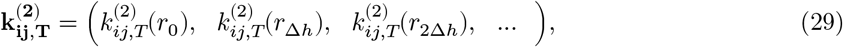

for some displacement step size Δ*h*, where each vector element 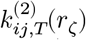 describes the density of (type *i*)-(type *j*) cell pairs that are separated by a distance *r* ∈ [*r*_*ζ*_, *r*_*ζ*_ + Δ*h*). Following Eq.8b, static secondorder spatial cumulants can be approximated by subtracting densities (Eq.27) from pair-densities (Eq.28). Then, analogously to Eq.29, following Eq.1, static second-order spatial cumulants can, in translationally invariant cases, be compactly approximated by a vector

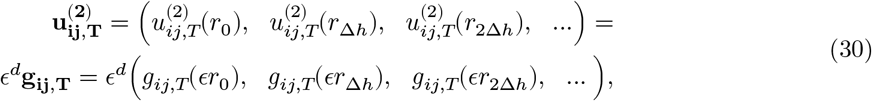

where the error terms *o*(*ϵ*^*d*^) have been omitted as they are set to zero in practical applications. In the computational pipeline codes developed for this study, we use the Fast Fourier Transform to calculate the integral in Eq.28, and thus approximate the spatial covariance (Eq.30). Note that the correction terms in Eq.1 generally depends on all vector elements in **g**_**ij**,**T**_.

### 2.10 Computer software and code

The `UF-Software` software is freely available to download (Cornell et al., 2019). The software comprises two main components: the `ModelSimulator` and the `ModelConstructor` which, respectively, can be used to generate STPPs and SCMs from user-defined RCP-process model descriptions. Note that, as MFPMs are a reduced form of SCMs, the `ModelConstructor` can also generate MFPMs. The `ModelSimulator` uses the Gillespie algorithm (Gillespie, 1977) and is written in the programming language C. The `ModelConstructor` is written in the Wolfram Mathematica software. Instructions on how to download the `UF-Software` are provided in the Supplementary Information (SI.1).

## 3 Methods and Model

### 3.1 The model implementation pipeline

For this study, we developed a model implementation pipeline (and pipeline codes) that systematises the use of the `UF-Software` in a number of steps (A1, B1-B4, C1-C7, D1 in Fig. 1). The pipeline includes a model formulation using RCP-processes (A1), the implementation of STPPs (B1-B4) and SCMs (C1C7), a visual comparison of STPP, SCM and MFPM-generated population dynamics (C7), and analysis of SCM and MFPM equations (D1). Note that, as MFPM equations are a reduced form of SCM equations, they are also formulated and calculated in the pipeline. Note also that the initial cell configurations (B2, C4) can be generated *in silico* or be obtained from biological data, and that spatial cumulants from point patterns from simulation data and biological data can be calculated (B4) and visualised (C3). If suitable for the application, analytical results from (D1) can be used in the model description. Instructions on how to access, run and modify the code files for each step in the pipeline are provided in the Supplementary Information (SI.1, SI.2).

**Fig. 1.**
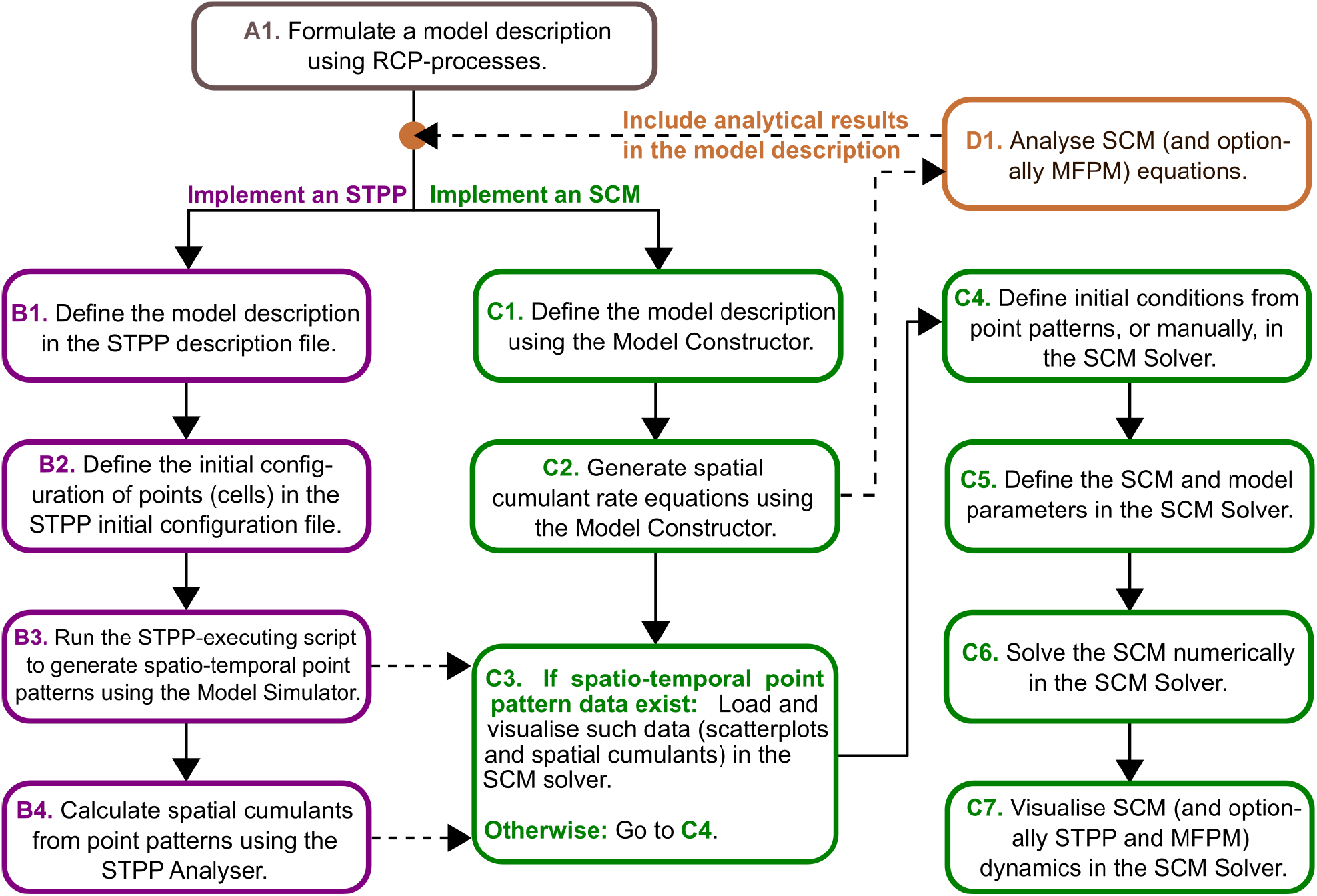
We developed a computational pipeline that streamlines the implementation, analysis and output comparison of spatio-temporal point processes (STPPs), mean-field population models (MFPMs) and spatial cumulant models (SCMs). The pipeline uses the `UF-Software` (Cornell et al., 2019) to generate STPPs (B1-B3), MFPMs and SCMs (C1-C2) from reactant-catalystproduct (RCP) process model descriptions (A1). STPP (B4) and SCM (C3-C7) generated summary statistics are thereafter calculated using the pipeline codes developed for this study. Note that, as MFPMs are reduced forms of SCMs, they are also handled in steps (C1-C7)

### 3.2 Describing a biological system using RCP-processes

The first step towards creating STPPs, SCMs and MFPMs using the computational pipeline entails translating a verbal model formulation into an RCP-formulation (Fig. 1, A1). In this study, as proof-of-concept, we consider a theoretical biological system that can be verbally described as follows:

*“Two types of cancer cells, s*_1_ *and s*_2_, *co-exist. Cells of type s*_1_ *produce growth factors that are both internalised by the producer cell and secreted to the environment. Cells of type s*_2_ *do not produce growth factors and must receive growth factors from s*_1_ *cells in order to proliferate. Both s*_1_ *and s*_2_ *cells can internalise growth factors from (other) s*_1_ *cells but, because s*_2_ *cells waste no energy on growth factor production, they benefit more from growth factor uptake than do s*_1_ *cells. Consequently, s*_2_ *cells have a higher growth factor-mediated proliferation rate than do s*_1_ *cells. Similarly, s*_1_ *cells express a higher proliferation rate when they receive growth factors from other cells, compared to when they self-produce growth factors. If drugs are applied to the system, s*_1_ *and s*_2_ *cells die at rates δ*_1_ *and δ*_2_, *respectively*.*”*

The ways in which cells can interact (and self-interact via autocrine signalling) through growth factors are pictorially shown in Fig. 2a-c, and the key conceptual difference between modelling STPP, SCM and MFPM interactions is pictorially described in Fig. 2d,e. The modelled system can be described using the RCP-processes listed in Table 1, and the default model parameter values are listed in Table 2.

**Fig. 2.**
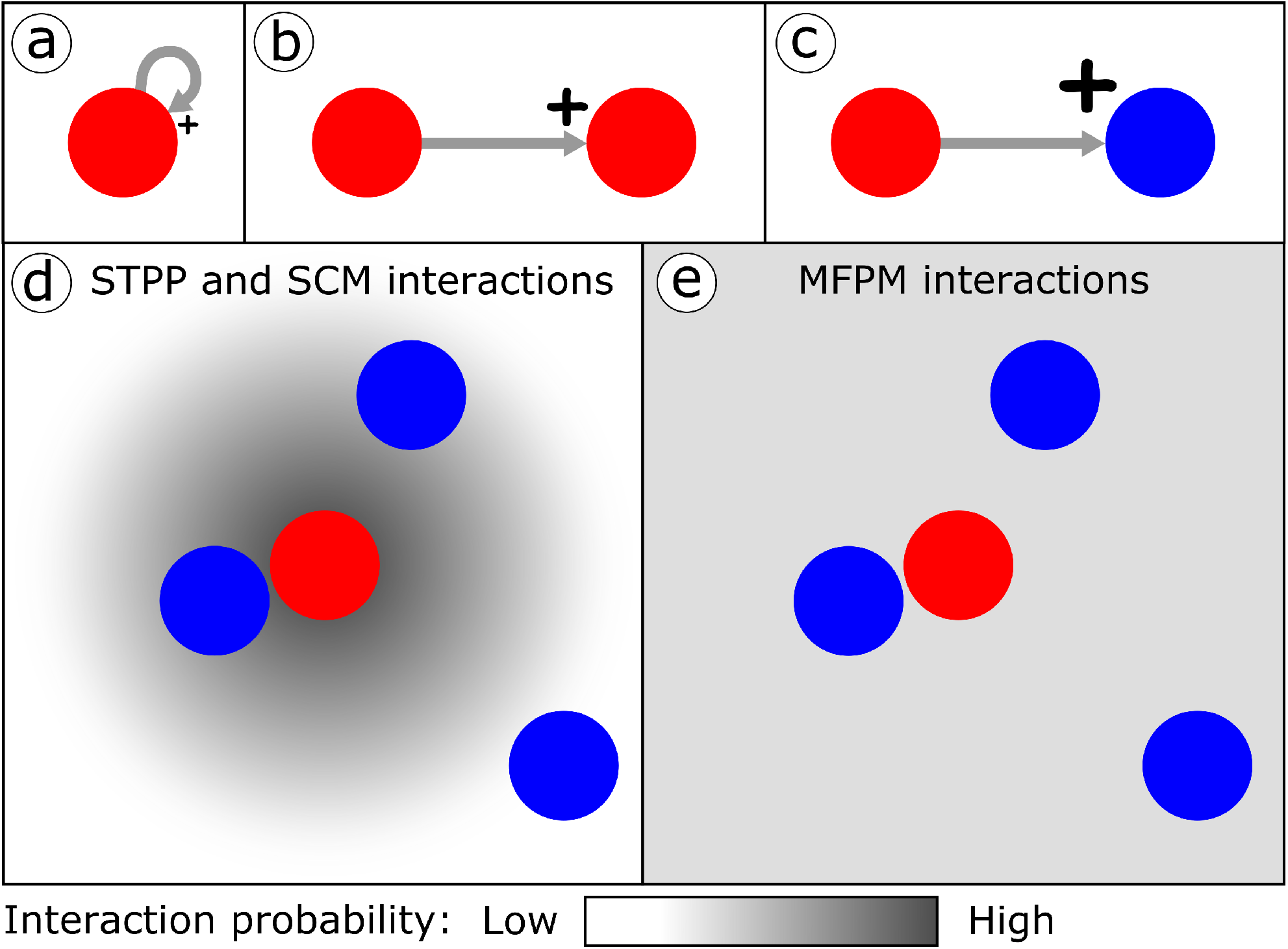
We consider a theoretical system in which cells interact with each other through growth factor secretion and uptake. Panels (a-c) show the possible ways in which cells in the modelled system can interact (and self-interact via autocrine signalling). The size of the plus (+) mark indicates the benefit (in terms of proliferation rate) that the receiver cell gains from the interaction. (a) Type *s*_1_ cells (red) can produce and internalise their own growth factors. (b) Type *s*_1_ cells can receive growth factors from other *s*_1_ cells. (c) Type *s*_2_ cells (blue) can receive growth factors from *s*_1_ cells. Panels (d,e) highlight the difference between modelling interactions in spatio-temporal point processes (STPPs), spatial cumulant models (SCMs) and mean-field population models (MFPMs). These panels show how likely the *s*_2_ cells are to receive growth factors from the focal *s*_1_ cell. In STPPs and SCMs, the probability that such a cell-cell interaction occurs depends on the distance between the two interacting cells, so that, for Gaussian interaction kernels, cells that are closer to each other are more likely to interact (d). In MFPMs, all cells in the system are equally likely to interact with each other, independently of spatial cell-cell separations (e)

**Table 1:** Biological processes are formulated in terms of reactant-catalyst-product (RCP) processes. The table includes the RCP-processes that are used to describe our theoretical system, in which growth factor producing (*s*_1_) and non-producing (*s*_2_) cells co-exist and interact.

| Biological process | Reactants (**r**),<br>catalysts (**c**),<br>products (**p**) | Markov<br>operator | Description |
| --- | --- | --- | --- |
| Birth$[s_1, B_1(x_1 - x_2)]$ | $\mathbf{r} = \emptyset$,<br>$\mathbf{c} = \{x_1, s_1\}$,<br>$\mathbf{p} = \{x_2, s_1\}$. | $L_B^{s_1}$ | A cell with mark $s_1$ in location $x_1$ produces a cell of mark $s_1$ in location $x_2$ with kernel $B_1(x_1 - x_2)$. |
| Birth By Facilitation<br>$[s_1, s_1, B_{11}(x_1 - x_2),$<br>$C(x_1 - x_3)]$ | $\mathbf{r} = \emptyset$,<br>$\mathbf{c} = \{\{x_1, s_1\}, \{x_3, s_1\}\}$,<br>$\mathbf{p} = \{x_2, s_1\}$. | $L_{BF}^{s_1 s_1}$ | A cell with mark $s_1$ in location $x_1$ produces a cell of mark $s_1$ in location $x_2$ with kernel $B_{11}(x_1 - x_2)$. This process is facilitated by a cell with mark $s_1$ in location $x_3$ with kernel $C(x_1 - x_3)$. |
| Birth By Facilitation<br>$[s_2, s_1, B_{12}(x_1 - x_2),$<br>$C(x_1 - x_3)]$ | $\mathbf{r} = \emptyset$,<br>$\mathbf{c} = \{\{x_1, s_2\}, \{x_3, s_1\}\}$,<br>$\mathbf{p} = \{x_2, s_2\}$. | $L_{BF}^{s_2 s_1}$ | A cell with mark $s_2$ in location $x_1$ produces a cell of mark $s_2$ in location $x_2$ with kernel $B_{12}(x_1 - x_2)$. This process is facilitated by a cell with mark $s_1$ in location $x_3$ with kernel $C(x_1 - x_3)$. |
| Density Independent<br>Death$[s_1, \delta_1]$ | $\mathbf{r} = \{x_1, s_1\}$,<br>$\mathbf{c} = \emptyset$,<br>$\mathbf{p} = \emptyset$. | $L_{DI}^{s_1}$ | A cell with mark $s_1$ in location $x_1$ dies at rate $\delta_1$. |
| Density Independent<br>Death$[s_2, \delta_2]$ | $\mathbf{r} = \{x_1, s_2\}$,<br>$\mathbf{c} = \emptyset$,<br>$\mathbf{p} = \emptyset$. | $L_{DI}^{s_2}$ | A cell with mark $s_2$ in location $x_1$ dies at rate $\delta_2$. |

**Table 2:**
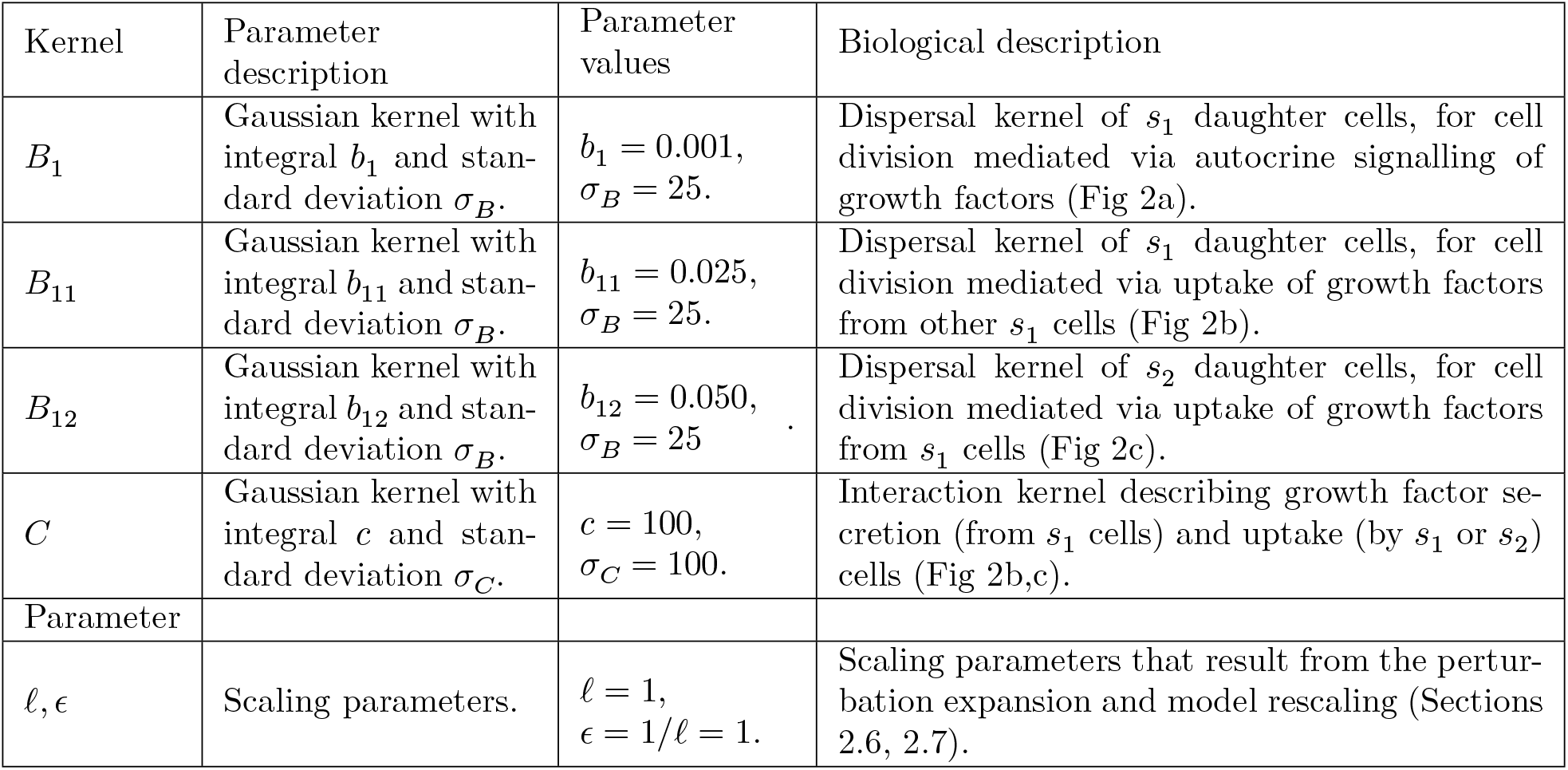
Model parameter values. The table shows the default model parameter values, in dimensionless form, that are used to produce the results in Figs. 3, 4, 5, 6.

| Kernel | Parameter<br>description | Parameter<br>values | Biological description |
| --- | --- | --- | --- |
| $B_1$ | Gaussian kernel with<br>integral $b_1$ and stan-<br>dard deviation $\sigma_B$. | $b_1 = 0.001$,<br>$\sigma_B = 25$. | Dispersal kernel of $s_1$ daughter cells, for cell<br>division mediated via autocrine signalling of<br>growth factors (Fig 2a). |
| $B_{11}$ | Gaussian kernel with<br>integral $b_{11}$ and stan-<br>dard deviation $\sigma_B$. | $b_{11} = 0.025$,<br>$\sigma_B = 25$. | Dispersal kernel of $s_1$ daughter cells, for cell<br>division mediated via uptake of growth factors<br>from other $s_1$ cells (Fig 2b). |
| $B_{12}$ | Gaussian kernel with<br>integral $b_{12}$ and stan-<br>dard deviation $\sigma_B$. | $b_{12} = 0.050$,<br>$\sigma_B = 25$. | Dispersal kernel of $s_2$ daughter cells, for cell<br>division mediated via uptake of growth factors<br>from $s_1$ cells (Fig 2c). |
| $C$ | Gaussian kernel with<br>integral $c$ and stan-<br>dard deviation $\sigma_C$. | $c = 100$,<br>$\sigma_C = 100$. | Interaction kernel describing growth factor se-<br>cretion (from $s_1$ cells) and uptake (by $s_1$ or $s_2$)<br>cells (Fig 2b,c). |
| Parameter |  |  |  |
| $\ell, \epsilon$ | Scaling parameters. | $\ell = 1$,<br>$\epsilon = 1/\ell = 1$. | Scaling parameters that result from the pertur-<br>bation expansion and model rescaling (Sections<br>2.6, 2.7). |

### 3.3 Describing a biological system using Markov operators

Each RCP-process corresponds to a Markov operator (Table 1), and an STPP can be described as a sum of such Markov operators. For our regarded model system, when no drugs are applied, the total Markov operator becomes

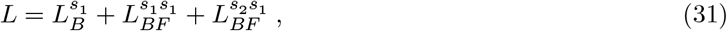

where all Gaussian birth kernels have the same standard deviations but different integrals such that

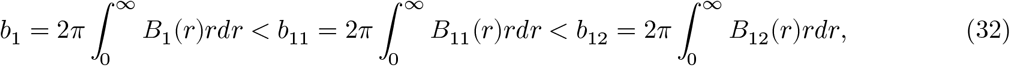

for *r* = |*x*_1_ − *x*_2_|, and the total integral of the cell-cell interaction kernel is

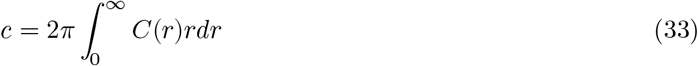

for *r* = |*x*_1_ − *x*_3_|. When drugs are applied to the system, the total Markov operator becomes

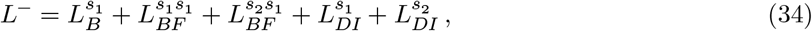

where the superscript (-) has been introduced to distinguish between the Markov operators in Eqs. 31 and 34. In this study, we set the parameter values pertaining to the Markov operators 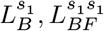 and 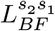 in step A1 (Fig. 1), and thus assume that these parameter values are inherent to the modelled cell population. Conversely, we assume that the parameters *δ*_1_ and *δ*_2_, pertaining to the Markov operators 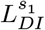 and 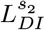 can be modulated by drug doses. Accordingly, in Section 4.2, we use MFPM and SCM equations to analytically derive death rates that result in non-growing subpopulations at an arbitrary treatment time *T* (Fig. 1, D1). These death rates are then implemented in the STPP, as implicitly modelled treatment responses.

### 3.4 Describing a biological system using MFPM and SCM equations

Using the `UF-Software`, rate equations for the variables *q*_*i*_(*t*), *p*_*i*_(*t*) and *g*_*ij*_(*t, ϵr*) in Eq.1 can be generated from RCP-formulated model descriptions. By implementing the model described by the Markov operator *L*^−^ (Eq.34) into the `UF-Software`, we obtain the following system of equations,

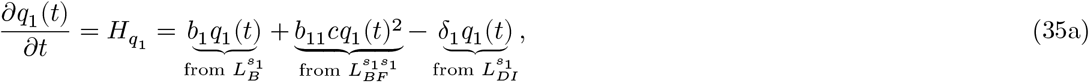

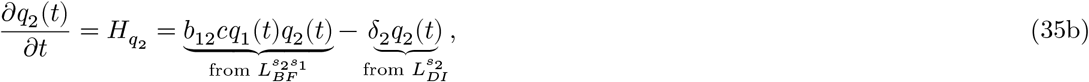

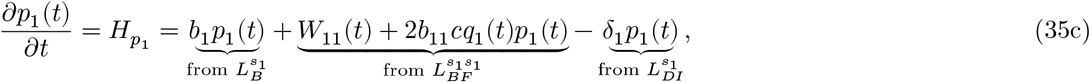

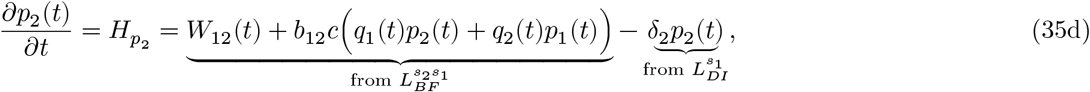

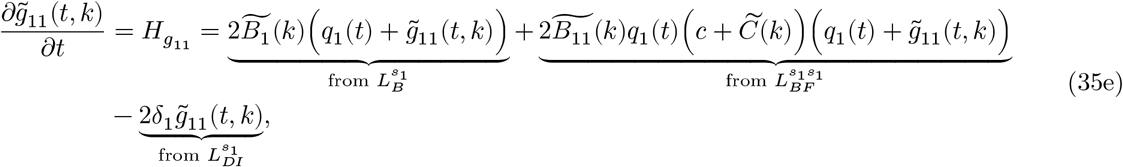

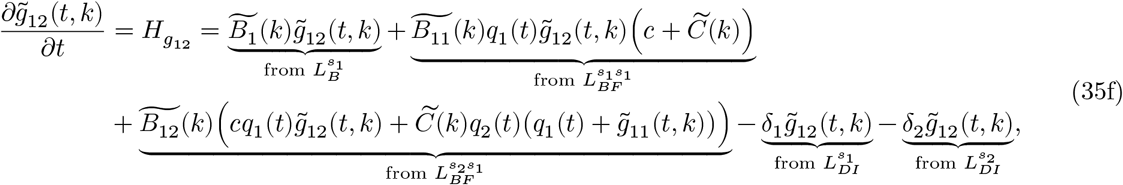

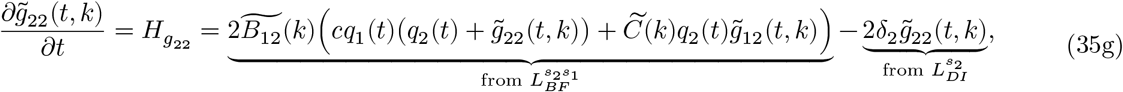

where 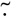 denotes the Fourier transform of ·, and *k* is the independent variable (the spatial frequency) in Fourier space. The *k*-notations have been introduced to denote the 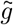-variables dependence on *k* and

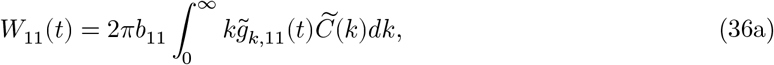

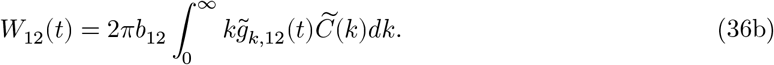

The SCM is described by Eqs.35(a-g) and Eqs.36a,b, whereas the MFPM is described by Eqs.35a,b only. Note that, in the absence of drugs, the terms resulting from the operators 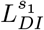 and 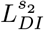 can be omitted so that the above system of equations describes the dynamics that corresponds to the Markov operator *L* (Eq.31). Equivalently, *δ*_1_ and *δ*_2_ can be set to zero.

### 3.5 Initial cell configurations and initial conditions

In this study, we consider three different initial cell configurations (ICCs). The ICCs have the same number of *s*_1_ and *s*_2_ cells, but different spatial configurations. Therefore, if we estimate the spatial cumulants from point patterns using the procedures outlined in Section 2.9, we find that the three ICCs have the same first-order spatial cumulants, but different second-order spatial cumulants (Fig. 3). In ICC.1, cells of type *s*_1_ and *s*_2_ are uniformly, randomly distributed across the simulated domain *D*. In ICC.2, cells of type *s*_2_ are uniformly, randomly distributed across *D*, whereas type *s*_1_ cells are randomly distributed within clusters. In ICC.3, both type *s*_1_ and *s*_2_ cells are clustered. In order to set the initial conditions for the SCM and MFPM models, we calculate densities and spatial covariances from the point patterns for all *i, j* = 1, 2. These calculations are used to set *q*_*i*_(*t* = 0) and *g*_*ij*_(*t* = 0, *ϵr*). The initial values for the SCM corrections are set to zero, so that *p*_*i*_(*t* = 0) = 0, for *i* = 1, 2.

**Fig. 3.**
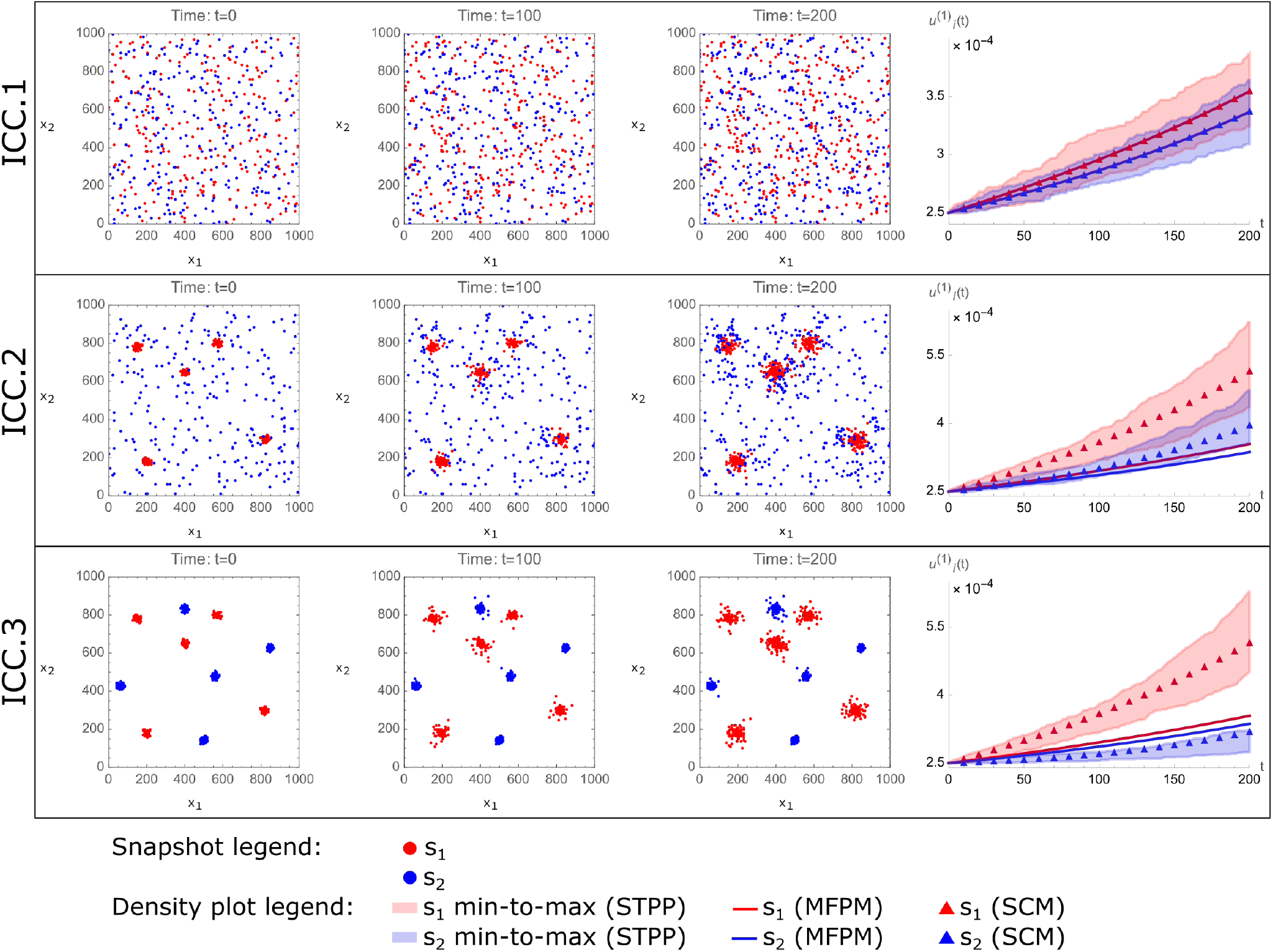
Spatial cumulant models (SCMs) capture density dynamics generated by spatiotemporal point processes (STPPs), even when mean-field population models (MFPMs) fail to do so. The figure shows STPP, MFPM and SCM-generated cell population dynamics, where each row corresponds to a unique ICC (left-most column). Snapshots from one STPP-generated spatio-temporal point pattern per ICC are shown at time points *t* = 0, *t* = 100, and *t* = 200. Growth factor-producing cells (type *s*_1_) are shown in red, and non-producers (type *s*_2_) are shown in blue. The right-most column shows STPP, MFPM and SCM-generated density dynamics. In the density plots, the shaded STPP data shows a span of minimum to maximum (min-to-max) density values calculated from 100 spatio-temporal point patterns resulting from the same ICC. MFPM (solid lines) and SCM (triangles) results are obtained by numerically solving the MFPM and SCM equations with initial conditions calculated from the shown ICCs. The parameter values used to create the plots are listed in Table 2

## 4 Results

### 4.1 SCMs capture STPP-generated cell population dynamics

We first set out to model the no-drug scenario with ICCs 1,2 and 3. To do this, we first implement the STPP (Eq.31) to generate 100 spatio-temporal point patterns from each ICC. We thereafter solve the MFPM (Eqs.35a,b) and SCM (Eqs.35a-g with Eqs.36a-b) equations numerically, using initial conditions for the spatial cumulants measured at the respective ICCs. In this proof-of-concept study, we will qualitatively say that a deterministic model “captures STPP dynamics”, if the model’s spatial cumulants lie within the minimum and maximum spatial cumulant values of 100 STPP-generated spatio-temporal point patterns.

Our results demonstrate that MFPM and SCM density dynamics coincide for ICC.1 only. For the simulated time duration, the MFPM is able to capture STPP-generated density dynamics resulting from ICC.1, where all cells are uniformly randomly distributed, but fail to do so for ICCs 2 and 3 (Fig. 3). As the cell-cell interactions via growth factors become more regional on the simulated domain, due to cell clustering, the MFPM performs worse in terms of describing STPP dynamics. This can be visually appreciated in Fig. 3, in which the deviation between MFPM and STPP dynamics increases for lower rows in the figure. On the other hand, the spatially informed SCM is able to capture STPP-generated density dynamics resulting from all considered ICCs in the simulated time duration (Fig. 3). This is due to the fact that SCMs account for spatio-temporal cell population structures by including equations that describe how the spatial covariances evolve in time. Indeed, our results show that spatial covariances calculated by SCMs capture those generated by STPPs (Fig. 4). In summary, the results in Figs. 3 and 4 show that SCMs can capture STPP-generated cell population dynamics, even when MFPMs fail to do so.

**Fig. 4.**
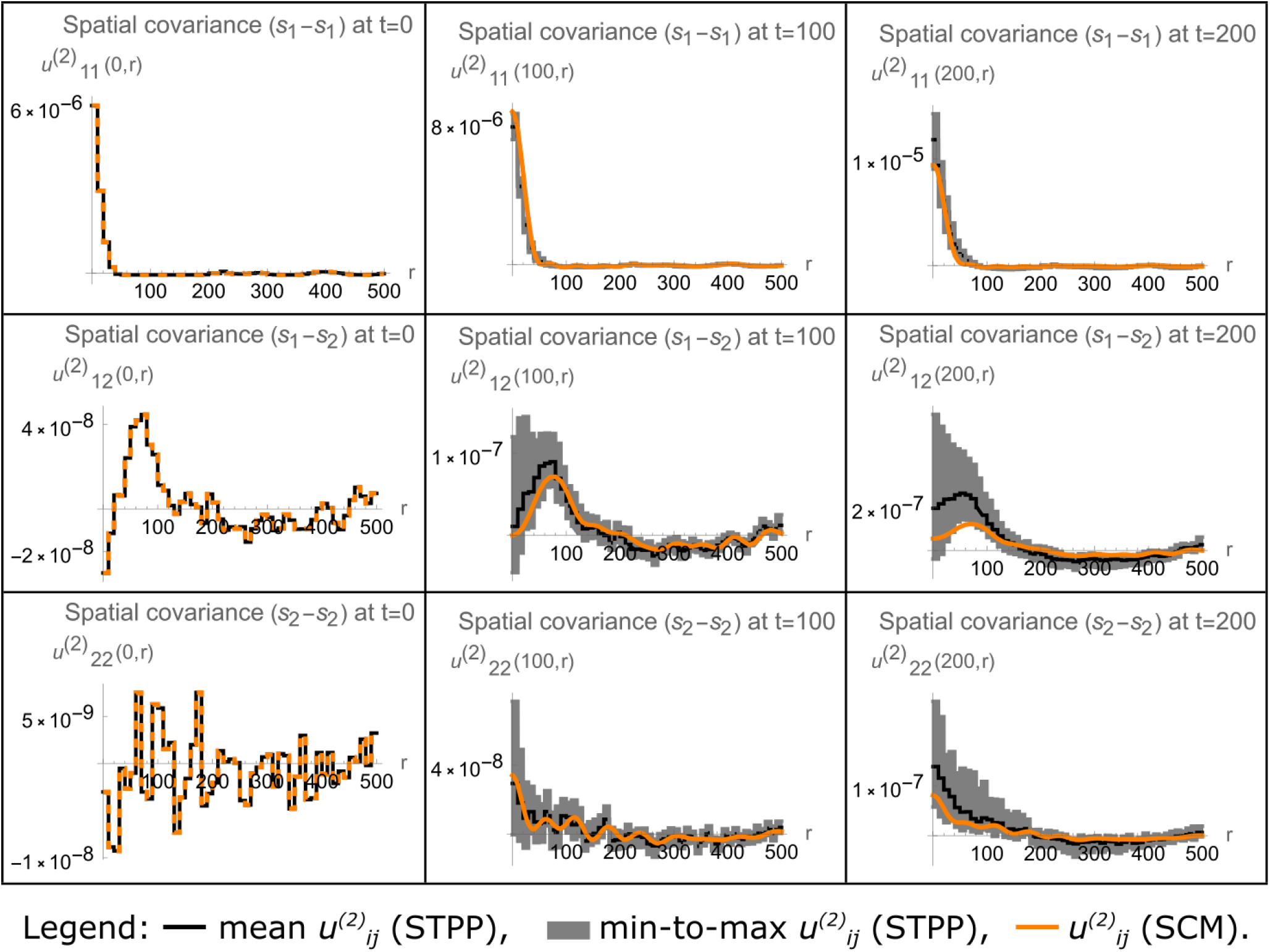
Spatial cumulant models (SCMs) capture spatial covariance dynamics generated by spatio-temporal point processes (STPPs). The figure shows STPP and SCM-generated spatial covariance dynamics resulting from ICC.2 (as shown in Fig. 3). Each row corresponds to a unique pairing of cell types, and each column corresponds to a time point (*t* = 0, *t* = 100, or *t* = 200.). Mean and minimum-to-maximum (min-to-max) values from 100 STPP-generated spatio-temporal point patterns are, respectively, shown with solid black lines and shaded gray bands. SCM results (orange) in the two right-most columns are obtained by numerically solving the SCM equations, with the spatial covariances shown in the left-most column as initial conditions. The parameter values used to create the plots are listed in Table 2

### 4.2 SCM-informed treatment strategies outperform MFPM-informed treatment strategies in STPP-generated cell populations

We next set out to derive theoretical drug doses, here implicitly modelled via cell death rates, that stop both subpopulations from growing. In other words, we want to find the model parameters *δ*_1_ and *δ*_2_ that cause the derivatives describing population growths to be zero at the treatment time *T*. We thus derive MFPM-informed drug doses by setting

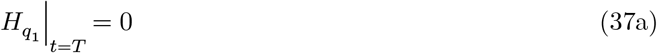

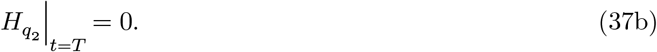

Solving Eqs.37a,b for 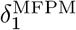 and 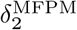, where the superscripts denote that these are the MFPMinformed doses, yields

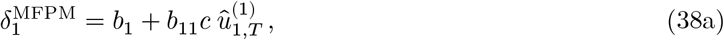

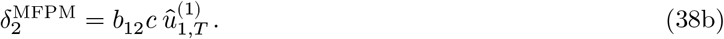

Here, the hat-notation 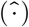 has been introduced to denote quantities that will be measured from point patterns, and the variables *q*_i_ (*t*) have been substituted with 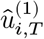, *i*.*e*., the measured density of subpopulation *i* at time *T*. Similarly, we derive SCM-informed drug by setting

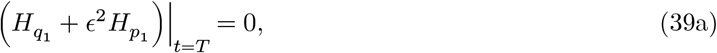

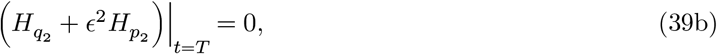

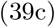

and solving for 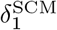 and 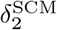 with all correction terms *p*_*i*_(*t*) set to zero upon measuring the densities and spatial covariances. Since all model parameters and variables are positive at time *T*, this yields

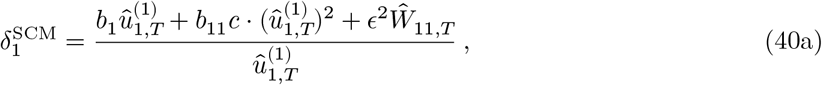

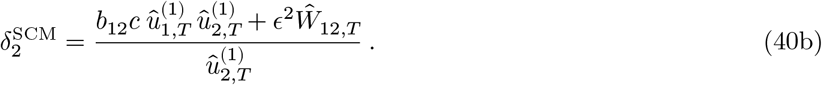

To test these drug doses in the STPP, we, for each spatio-temporal point-pattern, measure the quantities 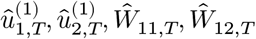, at the treatment time and calculate the MFPM and SCM-informed doses. Note that the quantities Ŵ_11,*T*_ and Ŵ_12,*T*_ incorporate the spatial structure of the regarded cell population (via spatial covariances) and local effects of growth factor cell-cell interactions (via the kernel *C*(*r*)). After time *T*, we then let each spatio-temporal point pattern progress further in time with its individual death rates *δ*_1_ and *δ*_2_ applied to the STPP (Eq.34).

Our results show that both MFPM and SCM-informed treatment doses are able to stop subpopulations *s*_1_ and *s*_2_ from increasing for cell populations that started at ICC.1 (Fig. 5, top row). However, for populations that started with ICC.2, the SCM-informed doses drastically outperform MFPM-informed doses in terms of inhibiting population growths (Fig. 5, middle row). In this case, the MFPM-informed doses underestimate the required values of *δ*_1_ and *δ*_2_ due to the synergistic growth effects that arise from cell type *s*_1_ − *s*_1_ proximity, and cell type *s*_2_ being close enough to *s*_1_ cells to benefit from their secreted growth factors. In cell populations initiated with ICC.3, type *s*_1_ cells are benefiting from being close to each other, whereas type *s*_2_ cells are too far away the from *s*_1_ cells to benefit from high *s*_1_ densities. Accordingly, the MFPM-informed doses adequately inhibit population growths for *s*_2_ but not for *s*_1_ (Fig. 5 bottom row). However, the SCM-informed doses inhibit both *s*_1_ and *s*_2_ population growths for ICC.3. In conclusion, SCMs enable the derivation of treatment strategies that are informed by (1) the spatial structure of the targeted cell population, and (2) by localised cell-cell interactions. In our theoretical study, and in terms of inhibiting subpopulation growths, SCM-informed doses match MFPM-informed doses for ICC.1, and outperform MFPM-informed doses for ICCs 2 and 3.

**Fig. 5.**
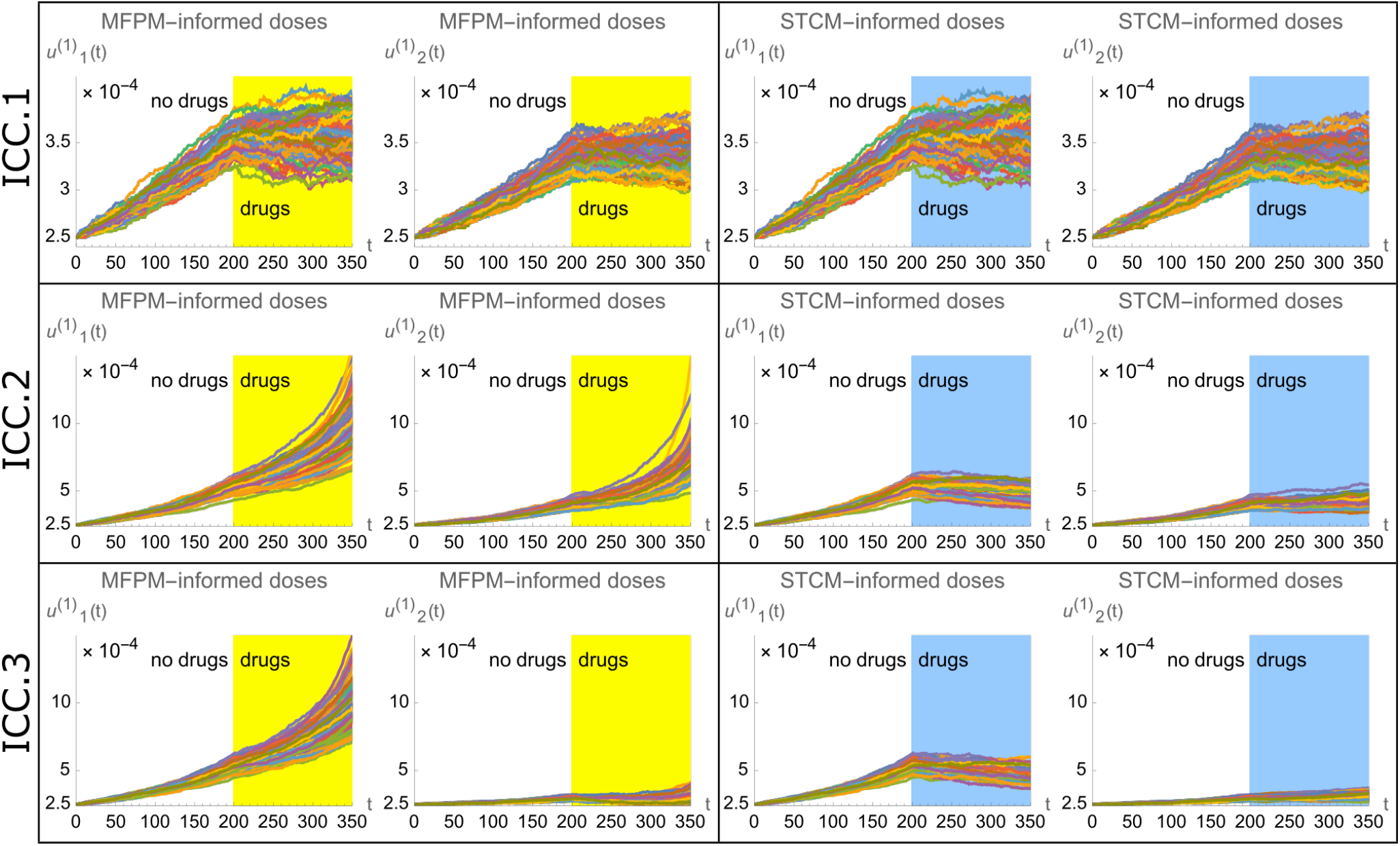
Treatment strategies informed by spatial cumulant models (SCMs) outperform treatment strategies informed by mean-field population models (MFPMs) in terms of inhibiting population growths in cell populations generated by spatio-temporal point processes (STPPs). The figure shows STPP-generated density dynamics when drugs are applied at *t* = 200. Each row corresponds to the dynamics resulting from a unique ICC (as shown in Fig. 3). Each panel includes two plots, one of which shows the density dynamics of the growth factor-producing cells 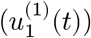, and one of which shows the density dynamics of the non-producer cells 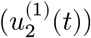. All plots show density dynamics from 100 spatio-temporal point patterns. Every spatio-temporal point pattern is assigned point pattern-specific drug doses (cell death rates) at *t* = 200, and is visualised with an individual line. Dynamics resulting from MFPM-informed drug doses (indicated by yellow areas) and SCM-informed drug doses (indicated by blue areas) are respectively shown in the left and right panels. The parameter values used to create the plots are listed in Table 2

### 4.3 Cell-cell interaction ranges and cell motility impact consistencies between STPP, MFPM and SCM density dynamics

Recall from Eq.1 that SCM subpopulation densities are expressed as MFPM densities plus a correction term. This correction term depends on the scaling factor *ϵ* = 1/ℓ, the densities and spatial covariances of the modelled population, and the RCP-processes that describe the actions and interactions of the modelled individuals via the variables *q*_*i*_(*t*), *p*_*i*_(*t*), *g*_*ij*_(*t, ϵr*). Consistencies (and deviations) between MFPM and SCM density dynamics thus depend both on the modeller’s choice of ℓ, and on properties of the modelled population. To demonstrate these dependencies, we revisit the model system studied in Section 4.1 (Fig. 3), and include cell motility in the system. We then vary two system properties, specifically the range of cell-cell interactions and the magnitude of cell motility, to investigate how well MFPMs and SCMs capture STPP-generated density dynamics in response to these variations. In order to include cell motility to the model system, we first introduce two new RCP-processes in Table 3 that describe random movement of type *s*_1_ and *s*_2_ cells. The model parameter values that are associated with these RCP-processes are listed in Table 4.

**Table 3:** Cell motility processes are formulated in terms of reactant-catalyst-product (RCP) processes. The table includes the RCP-processes that are used to describe random cell movement of growth factor-producing (*s*_1_) and non-producing (*s*_2_) cells.

| Biological process | Reactants (**r**),<br>catalysts (**c**),<br>products (**p**) | Markov<br>operator | Description |
| --- | --- | --- | --- |
| $\text{Jump}[s_1, M_1(x_1 - x_2)]$ | $\mathbf{r} = \{x_1, s_1\},$<br>$\mathbf{c} = \emptyset,$<br>$\mathbf{p} = \{x_2, s_1\}.$ | $L_M^{s_1}$ | A cell with mark $s_1$ in location $x_1$ jumps to location $x_2$ with kernel $M_1(x_1 - x_2)$. |
| $\text{Jump}[s_2, M_2(x_1 - x_2)]$ | $\mathbf{r} = \{x_1, s_2\},$<br>$\mathbf{c} = \emptyset,$<br>$\mathbf{p} = \{x_2, s_2\}.$ | $L_M^{s_2}$ | A cell with mark $s_2$ in location $x_1$ jumps to location $x_2$ with kernel $M_2(x_1 - x_2)$. |

**Table 4:**
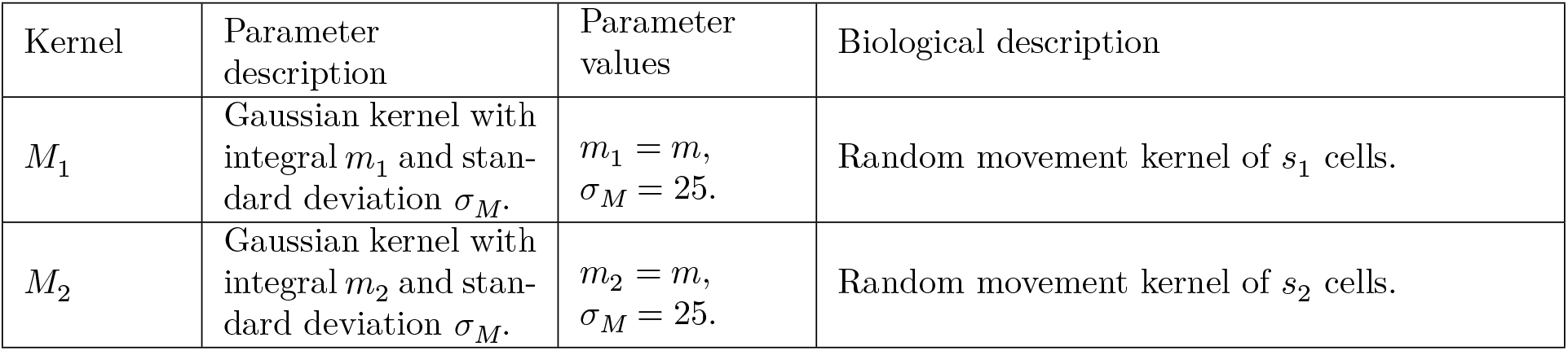
Model parameter values for random cell movement. The table shows the model parameter values, in dimensionless form, that are used to produce the results in Fig. 6. Note that the value of *m* is set on an example-specific basis (Fig. 6).

| Kernel | Parameter<br>description | Parameter<br>values | Biological description |
| --- | --- | --- | --- |
| $M_1$ | Gaussian kernel with<br>integral $m_1$ and stan-<br>dard deviation $\sigma_M$. | $m_1 = m,$<br>$\sigma_M = 25.$ | Random movement kernel of $s_1$ cells. |
| $M_2$ | Gaussian kernel with<br>integral $m_2$ and stan-<br>dard deviation $\sigma_M$. | $m_2 = m,$<br>$\sigma_M = 25.$ | Random movement kernel of $s_2$ cells. |

By adding the Markov operators listed in Table 3 to the Markov operator *L* in Eq.31, we formulate a new Markov operator *L*^+^ that describes a system in which growth factor-mediated cell division and random cell movement occur so that

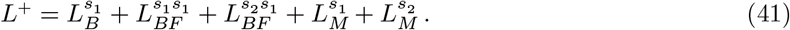

In order to generate the MFPM and SCM equations that correspond to Eq.41, note first that cell motility does not contribute to the MFPM equations. Therefore the corresponding MFPM model equations are Eqs.35a,b without cell death, so that the death rates are *δ*_1_ = 0 and *δ*_2_ = 0. Similarly, cell motility only impacts the SCM correction equations implicitly, via their dependence on the spatial covariance variables, but not explicitly. Therefore the SCM correction terms that correspond to the Markov operator *L*^+^ (Eq.41) are simply Eqs.35c,d with *δ*_1_ = 0 and *δ*_2_ = 0. The SCM rate equations for the spatial covariance variables do, however, need to be explicitly updated (from 35e-g) to include cell movement. We let the plus-notation (+) denote these updated rate equations and get

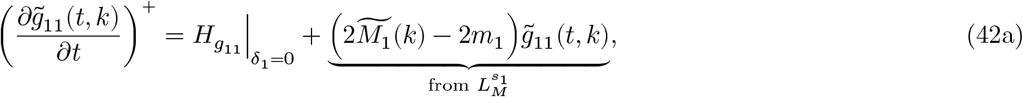

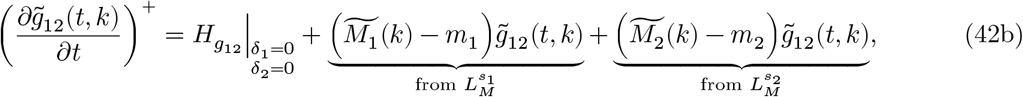

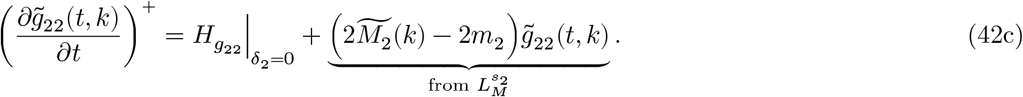

The terms 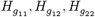 depend on spatio-temporal system dynamics and are listed in Eqs.35e-g. The Fourier transform of · is, again, denoted by 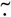 and *k* is the independent variable in Fourier space. In summary, the SCM equations that describe the population dynamics generated by an STPP that is governed by the Markov operator *L*^+^ (Eq. 41) are Eqs.35a-d (with *δ*_1_ = 0 and *δ*_2_ = 0), Eqs.36a,b, and Eqs.42a-c.

Now that the MFPM and SCM equations that correspond to *L*^+^ (Eq.41) have been formulated, we are ready to compare STPP, MFPM and SCM system dynamics for variations of the kernels *C* (Table 2) *M*_1_ and *M*_2_ (Table 4). In order for this example-specific comparison to yield qualitatively distinct outcomes, we use initial cell configuration ICC.3 in which the cells are strongly clustered (Fig. 3). Let us first consider the model system in which the cells are sessile, so that *m*=0 (Table 4), and vary *σ*_*C*_, the standard deviation of the cell-cell interaction kernel *C* that governs growth factor secretion and uptake (Table 2). In Fig. 6a, we compare STPP, MFPM and SCM density dynamics for three different values of *σ*_*C*_ that we classify to yield strongly localised (*σ*_*C*_ = 100), moderately localised (*σ*_*C*_ = 150) and weakly localised (*σ*_*C*_ = 200) cell-cell interactions.

**Fig. 6.**
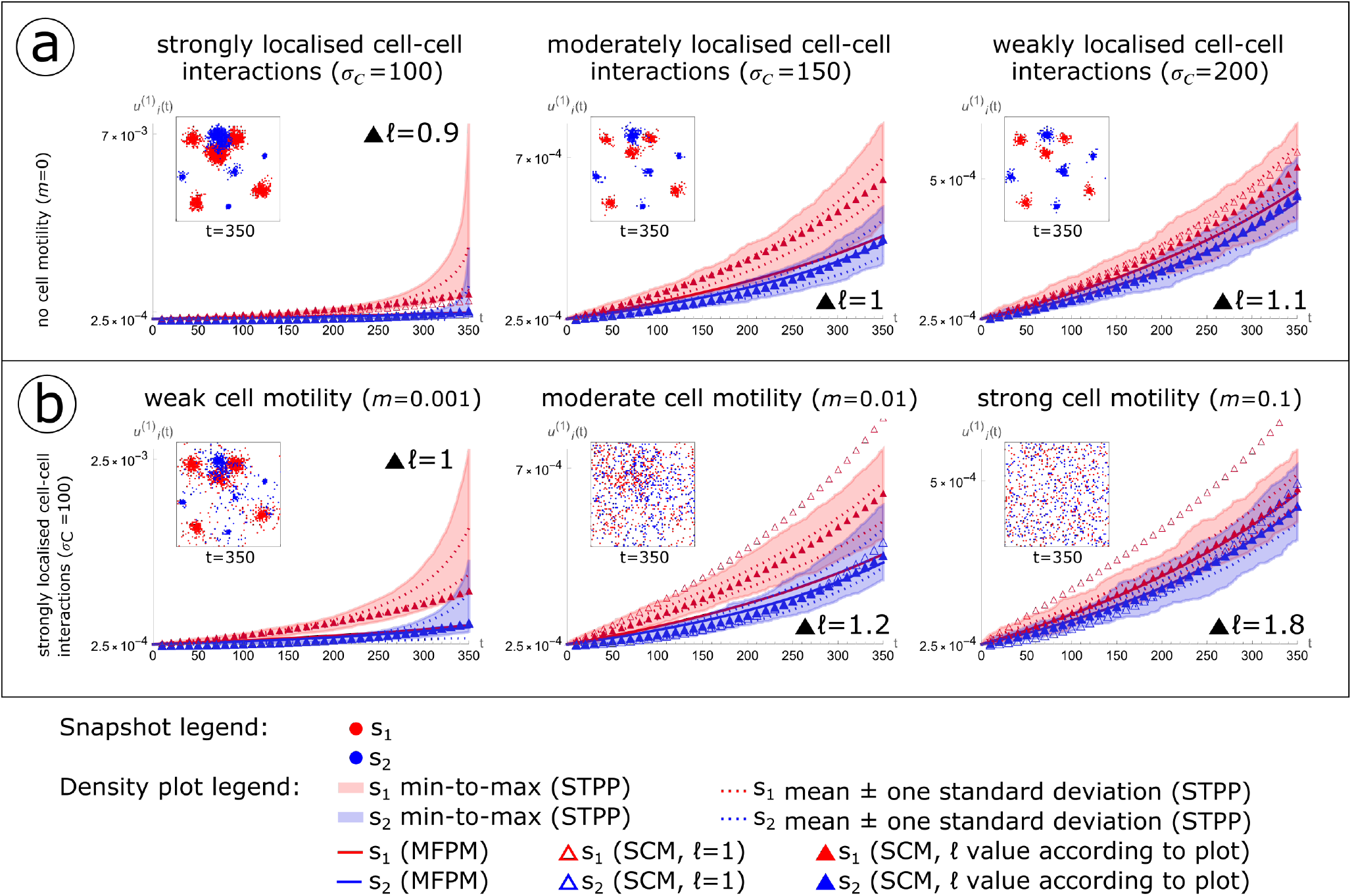
Cell-cell interaction ranges and cell motility impact consistencies between density dynamics generated by spatio-temporal point processes (STPPs), mean-field population models (MFPMs) and spatial cumulant models (SCMs). The figure shows STPP, MFPM and SCM cell population density dynamics resulting from the initial cell configuration ICC.3 shown in Fig. 3. (a) The standard deviation *σ*_*C*_ in the Gaussian kernel that governs cell-cell interactions via growth factors (*C* in Table 2) is different in each column. (b) The integral *m* of the Gaussian kernel that governs random cell movement is different in each column. In both (a) and (b), the inserted scatter plots show one STPP-generated spatio-temopral point pattern at the simulation end time *t* = 350. Growth factorproducing cells (type *s*_1_) are shown in red, and non-producers (type *s*_2_) are shown in blue. In the density plots, the shaded STPP data shows the span of minimum to maximum (min-to-max) density values from 100 spatio-temporal point patterns. Density mean values plus/minus one standard deviation from these spatio-temporal point patterns are shown with dotted lines. Deterministic MFPM (solid lines) and SCM (triangles) results are obtained by numerically solving the MFPM and SCM equations with initial conditions calculated from ICC.3. Unfilled triangles show SCM dynamics when ℓ = 1 (where the scaling parameter *ϵ* = 1/ℓ), and filled triangles show SCM dynamics when ℓ has been chosen according to the plots. Other parameter values used to create the plots are listed in Tables 2 and 4

We next consider the model system with a fixed *σ*_*C*_ value (*σ*_*C*_ = 100), and vary *m*, the total integral of the interaction kernels *M*_1_ and *M*_2_ (Table 4) that govern random cell movements. In Fig. 6b, we compare STPP, MFPM and SCM-generated density dynamics for three different values of *m* that we classify to yield weak (*m* = 0.001), moderate (*m* = 0.01) and strong (*m* = 0.1) cell motility.

In summary, Fig. 6 provides an example-driven, qualitative three-part stratification (I-III) of MFPM and SCM performance, in terms of their ability to capture STPP-generated density dynamics: (I) Cases where both MFPMs and SCMs underperform due to strongly localised cell-cell interactions and no-to-weak cell motility (Fig. 6a,b, left column). (II) Cases where MFPMs underperform whilst SCMs perform well due to moderately localised cell-cell interactions and no cell motility, or strongly localised cell-cell interactions and moderate cell motility (Fig. 6a,b, middle column). (III) Cases where both MFPMs and SCMs perform well due to weakly localised cell-cell interactions or strong cell motility (Fig. 6a,b, right column). This stratification (I-III) emphasises that a modeller’s choice of modelling method should be influenced by the model system at hand, and that the choice of *ϵ* = 1/ℓ in applications should be influenced by how much the MFPM dynamics deviates from the STPP dynamics.

In our problem-specific examples for case (I), SCM densities lie within the minimum-to-maximum densities of 100 STPP simulations, but not within the mean plus/minus one standard deviation of the simulations. Although the SCMs perform better than the MFPMs, in terms of capturing STPP dynamics, they underestimate STPP densities towards the end of the simulation time where STPP population sizes have increased by up to a factor 28 (Fig. 6a, left column). As we have not included any crowding or competition effects in our model system, this gives rise to strongly localised effects in strongly clustered populations that cause the SCMs, as formulated in the limit of long-ranged interactions, to underperform. In order to analytically describe STPPs in such cases, a modeller might attempt to consider other limits in which to develop SCM equations, or to use other spatial moment or cumulant techniques in which higherorder correction terms are included or strongly localised interactions are accounted for. Alternatively, research question permitting, a modeller might choose to use simulation-based approaches only, without an analytical counterpart, to study the dynamics of STPP-generated cell populations.

On the other hand, in case (III), MFPMs are sufficient to capture STPP dynamics and thus the inclusion of SCM model equations provides an unnecessary complication that can be avoided. Note that, according to the mean-field assumptions that underlie MFPMs, each cell in the modelled system interacts with any other cell in the system with equal probability (Fig. 2e). It is, therefore, to be expected that the appropriateness of these assumptions, and by extension the MFPMs ability to capture STPP-generated population dynamics, improves with increasing cell-cell interaction ranges (Fig. 6a) and cell motility (Fig. 6b).

Between the edge cases (I) and (III), lies case (II) in which SCMs provide an accurate and non-superfluous methodology to describe (Figs. 3,4,6) and manipulate (Fig. 6) STPP-generated cell population dynamics. We argue that there exist multiple applications in cell biology for which case (II) is relevant, and the use of STPPs and SCMs are appropriate. For example, STPPs and SCMs can be used to study cell systems in which cells are sessile or weakly to moderately motile, *i*.*e*., not motile enough to warrant mean-field assumptions. SCMs can also be used to study systems in which cell-cell interactions are localised but extend nearest neighbour interactions. Such cell-cell interactions include localised competition for space or nutrients, and secretion and uptake of diffusible factors that can affect cell division, cell death and cell motility.

## 5 Discussion

Most current cancer treatments target genomic aberrations without consideration for eco-evolutionary tumour aspects that change over the course of treatment, such as cell-cell and cell-microenvironment interactions. The lack of understanding of such dynamical tumour aspects is one factor that may be hindering the development of efficient targeted treatment protocols (Gatenby et al., 2020; Wang and Deisboeck, 2019). However, clinical trials that are informed by mathematical modelling and take dynamical tumour processes into account have started to emerge (Zhang et al., 2017; Poels et al., 2021).

Although not applied in clinical settings, IBMs are widely used in mathematical oncology to study cancer eco-evolution in response to cell-cell interactions and treatments. The appeal of IBMs has increased with the resolution of state-of-the-art experimental and clinical cancer data revealing information on the single-cell level. With their spatio-temporal structure, IBMs, unlike MFPMs, have the ability to include both spatial information about how subpopulations are located in tumours, from spatial transcriptomics (Hunter et al., 2021), and temporal information about how genetic alterations in tumour cell subpopulations propagate over generations to evade treatments, from phylogenetic analysis (Schwartz and Schäffer, 2017). However, without an analytical mathematical framework in which to analyse IBMs, it is difficult to describe relationships between model variables and model parameters in a mechanistic way that lets us perturb the modeled system and better understand its inter-component dependencies. In the context of cancer, theoreticians, experimentalists and clinicians may be interested in perturbing cancer systems onto evolutionary trajectories that are favourable from treatment perspectives. Analytical mathematical models that account for treatment effects and cell-cell interactions can provide instructions for how to do so (Kaznatcheev et al., 2019).

In this article, we demonstrated that SCMs enable spatially informed treatment strategies and analysis of local interactions in cancer systems. Although the theoretical, biological systems that we modelled in this study were of a simple structure, only comprising up to five RCP-processes, the model systems could be expanded to include more biological processes. It is straightforward to use the `UF-Software` and the pipeline codes developed for this article to implement processes such as crowding effects, density-dependent cell death and cell-type conversions via cell-intrinsic or external factors. We remark that SCMs could be further improved to include more functionalities that are relevant in cell biology. For example, in this study, we categorised cells with discrete marks, so that each cell belonged to one of two subpopulations with set traits. However, in many cell biology scenarios, it would be more relevant to use continuous marks to describe *e*.*g*., growth factor production, cell motility, drug resistance and gene expression in other forms. Moreover, in this study, all model parameters were cell-intrinsic constants. To allow for non-linear effects between a cell’s proximity to other cells and its behaviour, the model parameters could be adjusted to be functions of *e*.*g*., time, space or continuous marks. We leave these suggested improvements to the SCM framework as future challenges. Another future challenge will be to develop methods to parameterise the SCMs with experimental and, ultimately, clinical data. We emphasise that even though the mathematical theory that underlies SCMs may appear mathematically challenging at first sight, implementing SCMs (using the `UF-Software` and the pipeline codes developed for this study) is not. In order to make the methods discussed in this article accessible to the research community, we have provided clear instructions for how to access, run and modify these code files in the Supplementary Information (SI.1, SI.2).

Multiple research groups have mathematically modelled the effect that growth factor-mediated cellcell interactions have on cancer progression and treatment outcomes. Notably, Zhang *et al*. are using an EGT model to inform treatment strategies in a clinical prostate cancer trial (2017). Their model aims to predict personalised treatment outcomes in response to drug schedules and assumed interactions amongst testosterone producing, dependent and independent cells (see also (Mistry, 2021)). Kaznatcheev *et al*. used an EGT model to show that both drugs and cancer-associated fibroblasts modulate cell-cell interactions in co-cultures of drug-sensitive and resistant non-small cell lung cancer cells (2019). Haridas *et al*. used a PDE model to quantify how interactions between melanoma cells and fibroblasts affect the spreading of cells in *in vitro* co-cultures, finding no conclusive evidence of any cell-cell interactions via diffusible substrates (2017). Gerlee *et al*. used an IBM and an IBM-derived ODE model to demonstrate that autocrine signaling suffices to cause strong and weak Allee effects in cancer, in line with experimental observations from patient-derived brain tumor cell lines (2022).

We argue that SCMs provide a framework in which to study similar cancer research problems, with the added benefit of easy-to-produce duality: via the `UF-Software`, it is straightforward to generate both STPP-simulations and analytical SCM expressions describing user-defined biological systems. SCMs and can be especially helpful in studies when the locality of cell-cell interactions, via *e*.*g*., diffusible factors or resource competition, can not be neglected. Moreover, we remark that there exist parallels between the SCM and EGT frameworks, which both are centered around interactions amongst individuals, and we theorise that the SCM methodology of incorporating space may facilitate the derivation of corrected, spatially resolved EGT replicator equations. One important contrast between EGT models and SCMs, however, lies in how the modelled interactions are described. In EGT models, the behaviours of cell populations in response to cell-cell interactions are quantified in terms of gain or loss in proliferative fitness (Farrokhian et al., 2022). On the other hand, SCMs build upon rule-based descriptions of how the individuals (cells) in the modelled system act and interact via RCP-processes. SCMs can therefore be used to define ensembles of allowable biological mechanisms, *i*.*e*., candidate models, which could be evaluated against *in vitro* data, in line with a bottom-up modelling approach (Hamis et al., 2019). We also anticipate that the opportunity to analytically derive spatially informed cancer treatment strategies, as enabled via SCMs, will inspire new theoretical and applied research ventures.

## Code access

Instructions on how to access, run and modify the code files used in this study are available in the Supplementary Information (SI.1, SI.2).

