## Supplementary Information SI 1,2,3 for "Spatial cumulant models enable spatially informed treatment strategies and analysis of local interactions in cancer systems"

**Part III: The online Mathematica Notebook** 4

#### Part I:
### The code setup

#### SI.1 Downloading code files

##### SI.1.1 Downloading and installing the Unified Framework (UF) Software

**When using the UF-Software, cite:** Cornell, S.J., Suprunenko, Y.F., Finkelshtein, D. et al. A unified framework for analysis of individual-based models in ecology and beyond. Nat Commun 10, 4716 (2019). https://doi.org/10.1038/s41467-019-12172-y.

The `UF-Software`, which was developed by Cornell *et al.* 2019 can be downloaded from https://doi.org/10.6084/m9.figshare.9633161. The software contains two main parts: the `Model Constructor` and the `Model Simulator`.

The `Model Constructor` is a Wolfram Mathematica toolbox that enables the generation of spatial cumulant model (SCM) and mean-field population model (MFPM) equations. The toolbox comprises five Mathematica packages and requires Mathematica version 10 or greater. After downloading the packages, the packages must be installed in order to be used. This can be done by locating the `File`-tab in the Mathematica menu-bar and thereafter selecting `Install`, and entering the following information in the fields, `Type of Item to Install:` `Package`, `Source:` (locate the package), `Install Name:` (enter the name of the package).

The `Model Simulator` is a C code that enables the generation of spatio-temporal point processes (STPPs). It requires a C compiler.

##### SI.1.2 Downloading the pipeline code files

The pipeline developed in this study utilises the `UF-Software`. Pipeline codes can be downloaded from the code-hosting platform GitHub (https://github.com/SJHamis/SpatialCumulantModels_CancerApplications).

**When using the pipeline codes, cite the main article**: Hamis, S., Somervuo, P., Ågren, J.A. et al. Spatial cumulant models enable spatially informed treatment strategies and analysis of local interactions in cancer systems. (2022).

##### SI.1.3 The structure of code file directories

In order for the pipeline codes to work, the files must be arranged as shown in Figure SI.1.1. The location of the directory containing the `Model Constructor` packages does not impact the code files.

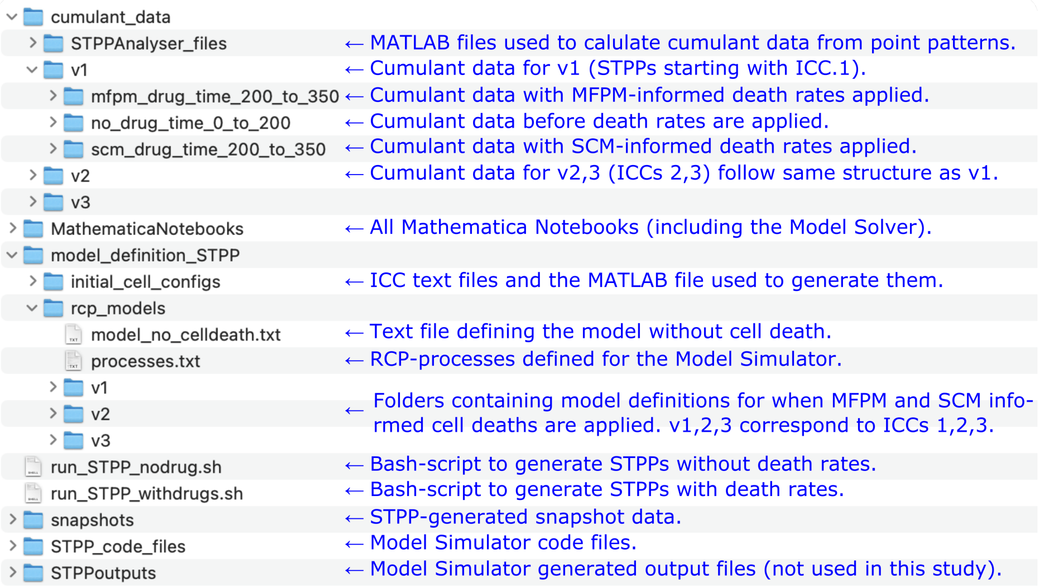

Figure SI.1.1: The structure of the pipeline code files, as embedded with the `UF-Software` code files.

#### Part II:
### The model implementation pipeline

Figure 1 in the main article provides a pictorial overview of the model implementation pipeline. The notation (A1,B1-4,C1-7,D1) used in this document follows that figure. The code files outlined in Sections SI.2-SI.2 describe the biological system presented in Section 3.2 of the main article. Detailed instructions on how to customise the code files to model other systems are provided in this document and via comments embedded in the code files.

#### SI.2 A: Model definition

##### SI.2.1 A1: Formulate a model description using RCP-processes

As a first step towards creating STPPs and deriving and SCM equations using the `UF-Software`, one must describe the modelled system by one or more reactant-catalyst-product (RCP) process (Cornell et al., 2019). Such model descriptions can, for clarity, be presented with the help of RCP-process tables. The `UF-Software` includes a set of pre-defined RCP-processes, some of which are included in Table SI.I. The software also allows a user to add custom RCP-processes (Cornell et al., 2019). A full biological system can described as a sum $L$ of Markov operators so that if we, for example, want to model a biological system in which cells with mark 1 divide, move and die (all in a density-independent way) then, using the tabulated Markov operators, $L = L_B^1 + L_J^1 + L_{DI}^1$.

| Biological process | Reactants ($\mathbf{r}$), catalysts ($\mathbf{c}$), products ($\mathbf{p}$) | Markov operator | Description |
|---|---|---|---|
| Birth$[i, a(x_1 - x_2)]$ | $\mathbf{r} = \varnothing$ , $\mathbf{c} = \{x_1, i\}$ , $\mathbf{p} = \{x_2, i\}$ . | $L_B^i$ | An individual with mark $i$ in location $x_1$ produces an individual of mark $i$ in location $x_2$ with kernel $a(x_1 - x_2)$. |
| Jump$[i, a(x_1 - x_2)]$ | $\mathbf{r} = \{x_1, i\}$ , $\mathbf{c} = \varnothing$ , $\mathbf{p} = \{x_2, i\}$ . | $L_J^i$ | An individual with mark $i$ jumps from location $x_1$ to $x_2$ with kernel $a(x_1 - x2)$. |
| Density Independent Death$[i, r]$ | $\mathbf{r} = \{x_1, i\}$ , $\mathbf{c} = \varnothing$ , $\mathbf{p} = \varnothing$ . | $L_{DI}^i$ | An individual with mark $i$ in location $x_1$ dies at rate $r$. |
| Death By External Factor$[i, j, a(x_1 - x_2)]$ | $\mathbf{r} = \{x_2, j\}$ , $\mathbf{c} = \{x_1, i\}$ , $\mathbf{p} = \varnothing$ . | $L_{DE}^{ij}$ | An individual of mark $i$ in location $x_1$ induces death in an individual of mark $j$ in location $x_2$ with kernel $a(x_1 - x_2)$. |

Table SI.I: A selection of pre-defined RCP-processes that are included in the `UF-Software`.

#### SI.2 B: Implement the spatio-temporal point process

##### SI.2.2 B1: Define the model description

The `Model Simulator` reads the RCP-processes model description from a text file. In the pipeline, this text file for the no-drug scenario is named `model_definition_STPP/model_no_celldeath.txt`. The file can be modified to describe other biological systems. The pre-defined RCP-processes that are included in the `UF-Software`, and the formats required to include them in model description text files, are listed in the file `model_definition_STPP/processes.txt`. The `UF-Software` recognises two types of kernels: Gaussian kernels and Top-Hat kernels, which are respectively written as `truncatedGaussian[`*`Integral, StandardDeviation`*`]` and `tophat[`*`Integral,Radius`*`]` in the model description text files, where *`Integral, StandardDeviation, Radius`* are numerical values.

##### SI.2.3 B2: Define the initial point configuration

In this study, we generated the initial cell configurations *in silico* using the MATLAB code `model_definition_STPP/initial_cell_configs/cerateInitialCellConfigurations.m`. The initial cell configuration data are written to the text files `model_definition_STPP/initial_cell_configs/ICC`*i*`.txt`, for $i$=1,2,3. In these text files, the first column denotes cell marks, and the other two columns denote cell coordinates. These text files can be modified to describe other initial cell configurations, which can be generated *in silico* or obtained be from biological data.

##### SI.2.4 B3: Generate spatio-temporal point patterns

In order to generate spatio-temporal point patterns from the STPP, a user can run the Bash-script `run_STPP_nodrug.sh` for the no-drug model from a terminal. In the Bash-scripts, variables can be edited to modify the model definition (`RCP_MODEL`), initial cell configuration (`ICC`), the number of spatio-temporal patterns to generate (indicated by the number of for-loops), the size of the spatial domain (for a quadratic domain, the number after option `-U` denotes the domain side), the simulation time (the number after option `-T`), the time intervals with which snapshot data will be taken (the number after option `-dT`). Note that the executable C file `STPP_code_files/ppsimulator` must be compiled for the Bash-script to work (in Linux systems this can be done by the command line `gcc ppsimulator.c -lm -O2 -o ppsimulator`), this only needs to be done once. For more information on how to customise the `ModelSimulator`, we refer the reader to Cornell *et al.*'s 2019 Supplementary Notes.

##### SI.2.5 B4: Evaluate point pattern data

The MATLAB file `cumulant_data/STPPAnalyser_files/measure_cumulants_from_pointpatterndata.m` can be run to generate approximate spatial cumulant data from point pattern data. More precisely, the file calculates mean, one standard deviation, minimum and maximum densities for all subpopulations at the sampled time points. The file also calculates mean, one standard deviation, minimum and maximum spatial covariances for all possible cell-mark pairs at user-defined time points (as a default, selected to be the initial, middle and final time points of the STPP simulations). These calculations utilise the Cross-Correlation Theorem. A user can modify the number of marks (*e.g.,* denoting cell types) in the MATLAB file by modifying the variable `no_subpops`. Output files are located in the folder `cumulant_data/vi`, where `i` corresponds to the ICC.

#### SI.2 C: Implement the spatial cumulant model

##### SI.2.6 C1-C2: Define the model description and generate SCM equations

By defining application-specific model description in the Mathematica Notebook `MathematicaNotebooks/generateODEs.nb`, SCM (and thus also MFPM) equations can be generated. Instructions on how to modify model descriptions are provided in the Notebook.

##### SI.2.7 C3-C7: Calculate SCM dynamics and compare with STPP and MFPM dynamics

The Mathematica Notebook `MathematicaNotebooks/SCMSolver.nb` can be used to (1) visualise snapshot data, (2) visualise spatial cumulant data, (3) calculate SCM and MFPM dynamics and (4) generate plots to visually compare STPP, SCM and MFPM dynamics. This Notebook produces the plots shown in Figure 3 and 4 in the main article. Instructions on how to modify the code to describe other user-defined biological systems are provided in the Notebook.

#### SI.2 D: SCM analysis

##### SI.2.8 D1: Analyse SCM and MFPM equations

The Mathematica Notebook `MathematicaNotebooks/deriveDeathRateExpressions.nb` is used to symbolically derive the MFPM and SCM informed death rates.

##### SI.2.9 Include analytical results in the model description

The symbolic death rate expressions calculated in the Notebook `MathematicaNotebooks/deriveDeathRateExpressions.nb` are functions of spatial cumulants, which generally will be different for each spatio-temporal point pattern. In order to find numerical death date values for each spatio-temporal point pattern, the spatial cumulants for each spatio-temporal point pattern at the treatment time are used. The death rates are calculated the Mathematica Notebook `MathematicaNotebooks/calculatePPSpecficDeathRates.nb`, which produces model description text files for both MFPM and SCM informed doses. These output files are respectively saved as `model_definition_STPP/rcp_models/`*vi*`/Model_MFPM_j` and `model_definition_STPP/rcp _models /`*vi*`/Model_SCM_j`. Here, `v`*i* is `v1` or `v2` or `v3`, where *i* denotes which initial cell configuration was used in the STPP. The spatio-temporal point patterns are labeled by *j*, where *j*`=1,2,..,S` and `S` denotes the number of spatio-temporal point patterns.

STPPs using spatio-temporal point pattern-specific death rates can be implemented using the Bash-script `run_STPP_withdrugs`. This generates post-drug snapshot (point pattern) data in the folder `snapshots/vi_dMFPM/runj` and `snapshots/vi_dSCM/runj`, using the labels `MFPM/SCM, i, j` as described in the above paragraph.

Single-track density data pre and post drug treatments can be generated by the MATLAB file `cumulant_data/STPPAnalyser_files/createSingleTrackDensityData.m`. One track correspond to one spatio-temporal point pattern. A user can modify the number of marks (*e.g.,* denoting cell types) in the MATLAB file by modifying the variable `no_subpops`. Output files are saved in the folders `cumulant_data/STPPAnalyser_files/vi`, for `i`=1,2,3.

The Mathematica Notebook `MathematicaNotebooks/evaluateModelDynamicsWithDrugs.nb` can be used to generate single-track density plots, pre and post drug treatments, as shown in Figure 5 in the main article.

#### Part III:
### The online Mathematica Notebook

The SCM methods used in our study are summarised in an online Mathematica Notebook on the Wolfram Community website (https://community.wolfram.com/groups/-/m/t/2559644). This Notebook provides an accessible overview of how we formulated, calculated and analysed SCMs in this study. Note that the online Notebook is a reduced version of the study's GitHub directory (Fig. SI.1.1).
